# Amine recognizing domain in diverse receptors from bacteria and archaea evolved from the universal amino acid sensor

**DOI:** 10.1101/2023.04.06.535858

**Authors:** Jean Paul Cerna-Vargas, Vadim M. Gumerov, Tino Krell, Igor B. Zhulin

## Abstract

Bacteria contain many different receptor families that sense different signals permitting an optimal adaptation to the environment. A major limitation in microbiology is the lack of information on the signal molecules that activate receptors. Due to a significant sequence divergence, the signal recognized by sensor domains is only poorly reflected in overall sequence identity. Biogenic amines are of central physiological relevance for microorganisms and serve for example as substrates for aerobic and anaerobic growth, neurotransmitters or osmoprotectants. Based on protein structural information and sequence analysis, we report here the identification of a sequence motif that is specific for amine-sensing dCache sensor domains (dCache_1AM). These domains were identified in more than 13,000 proteins from 8,000 bacterial and archaeal species. dCache_1AM containing receptors were identified in all major receptor families including sensor kinases, chemoreceptors, receptors involved in second messenger homeostasis and Ser/Thr phosphatases. The screening of compound libraries and microcalorimetric titrations of selected dCache_1AM domains confirmed their capacity to specifically bind amines. Mutants in the amine binding motif or domains that contain a single mismatch in the binding motif, had either no or a largely reduced affinity for amines, illustrating the specificity of this motif. We demonstrate that the dCache_1AM domain has evolved from the universal amino acid sensing domain, providing novel insight into receptor evolution. Our approach enables precise “wet”-lab experiments to define the function of regulatory systems and thus holds a strong promise to address an important bottleneck in microbiology: the identification of signals that stimulate numerous receptors.

## Introduction

Bacteria have evolved numerous receptors that sense environmental stimuli and modulate various signal transduction pathways to enable adaptation to changing conditions. Major receptor families include transcriptional regulators, sensor histidine kinases, chemoreceptors, cyclic (di)nucleotide cyclases and phosphodiesterases, serine/threonine protein kinases and phosphatases (1, 2). The initial step of signal integration by these receptors involves ligand binding to sensor, or ligand binding domains (LBD). Hundreds of different sensor domains have evolved, although only a few of them are ubiquitous (1, 3), and the same type of a sensor domain is frequently found in different signal transduction systems (4). Signals that activate most of signal transduction systems in bacteria and archaea are unknown, which presents a major bottleneck in microbiological research (5). Such knowledge is indispensable not only for understanding the physiological significance of regulatory circuits, but also for the development of anti-infective therapies aimed at reducing bacterial virulence by interfering with signal transduction cascades. Revealing signals for thousands of unstudied receptors by extrapolation from a few well-characterized homologs is largely hampered by the fact that sensor domains are rapidly evolving thus displaying a large degree of sequence divergence (6).

We have recently reported the first study in which the type of signal molecules was successfully predicted and verified for a large sensor domain family. By combining sequence and structure information from a few known amino acid sensing chemoreceptors of the ubiquitous dCache_1 domains (7) we derived the amino acid recognizing motif, which was used in database searches to identify thousands of motif-containing homologs, followed by experimental validation of selected targets (8). Subsequently, we defined a large subfamily of amino acid-sensing dCache_1 domains (termed dCache_1AA) containing more than ten thousand protein sequences from all major lineages of life (8). This iterative computational and experimental approach has an enormous potential to link many thousands of receptors to specific ligands, which is crucial for understanding the function of corresponding signal transduction circuits.

dCache_1 domains are the predominant extracellular sensors found in all major receptor families in bacteria and archaea (7). In addition to binding amino acids, some dCache_1 domains bind organic acids (9, 10), sugars (11), quorum sensing signals (12), inorganic ions (13), purine derivatives (14), polyamines (15, 16) and quaternary amines (QA)(17, 18), suggesting that in addition to dCache_1AA there might be other domain subfamilies with a well-defined ligand repertoire.

In this study, we focused on biogenic amines, because of their important biological roles and the availability of three solved structures of amine-bound dCache_1 domains from bacterial receptors. Biogenic amines are products of amino acid metabolism and are characterized by a nitrogen atom that is covalently linked to two, three or four alkyl substituents, resulting in secondary, tertiary, or quaternary amines. This compound family is present throughout the Tree of Life and its members possess a diverse range of biological functions. For example, choline is required for membrane phospholipid synthesis (19), acetylcholine is the major neurotransmitter (20), glycine-betaine and carnitine are important osmo- and cryoprotectants (21, 22), trimethylamine N-oxide (TMAO) is an electron acceptor for anaerobic respiration in bacteria (23) and methyl-, dimethyl-, and trimethylamines are important growth substrates for methanogenic archaea in habitats ranging from marine environments (24) to the human gut (25).

Here, we identify a sequence motif for biogenic **<u>am</u>**ine binding in dCache_1 domains (termed dCache_1AM) and show that it evolved from the ubiquitous amino acid binding motif dCache_1AA, which is found throughout the Tree of Life. This study further demonstrates that our approach is applicable to characterizing other ligand-binding domain families thus leading to substantial gain in knowledge on signal transduction systems.

## Results

### Different orientation of ligands in quaternary amine binding dCache_1 domains

We analyzed several solved structures of dCache_1 sensory domains in complex with quaternary amines: McpX from *Sinorhizobium meliloti* in complex with proline betaine (PDB ID 1R9Q) (26), PacA from *Pectobacterium atrosepticum* in complex with betaine (trimethyl glycine; PDB ID 7PSG) (17), and PctD from *Pseudomonas aeruginosa* in complexes with choline (PDB ID 7PRQ) and acetylcholine (PDB ID 7PRR) (17). Although the ligands are found in the same binding pocket in all three chemoreceptors, they are oriented in various directions with respect to their oxygen and nitrogen atoms (**Fig. 1A-D**). This is in stark contrast with the dCache_1AA domains, where various amino acids bind in the same orientation (8). This phenomenon was observed not only between receptors, but also in the same receptor with different quaternary amine ligands (**Fig. 1AB**). However, we noticed a common feature for all bound ligands – the cation-π interaction with the π-system of aromatic residues in the ligand binding pocket. For example, in the PctD ligand binding pocket choline and acetylcholine are oriented in opposite directions, but both make cation-π bonds with Y206 and W155 (**Fig. 1AB**). In PacA, betaine interacts with Y186, also through the cation-π bond (**Fig. 1C**) and proline betaine in McpX interacts with the favorably positioned Y139 (**Fig. 1D**). Considering that bonding energy of the cation-π interaction can be significant (27, 28), this type of bond may be of special importance for quaternary amine binding, acting as a “hinge” around which ligands can be oriented in various directions. In addition, several residues that form the ligand binding interface and provide stabilization of bound ligands are conserved among the receptors (**Fig. 1**, residues in green). Specifically, proline betaine makes a weak hydrogen bond with D208 of McpX (**Fig. 1D**), betaine makes an ionic bond with the corresponding residue in PacA (**Fig. 1C**), while choline and acetylcholine make hydrogen bonds with the corresponding D235 in PctD (**Fig. 1A,B**). In both McpX and PacA quaternary amines also make two hydrogen bonds with two non-conserved residues (G181/E180 and S159/Q165, respectively) that “lift” the oxygen containing end of the ligands (See **Fig. 1CD**). Thus, it appears that all three quaternary amine binding dCache_1 domains (PctD, PacA, ad McpX) might share a conserved ligand-binding motif. Another unique feature of the three quaternary amine binding receptors is the presence of an additional alpha-helical element above the binding pocket, where two charged residues (R103 and D133 in PctD) interact via two hydrogen bonds (**Fig. 2A**).

**Figure 1.**
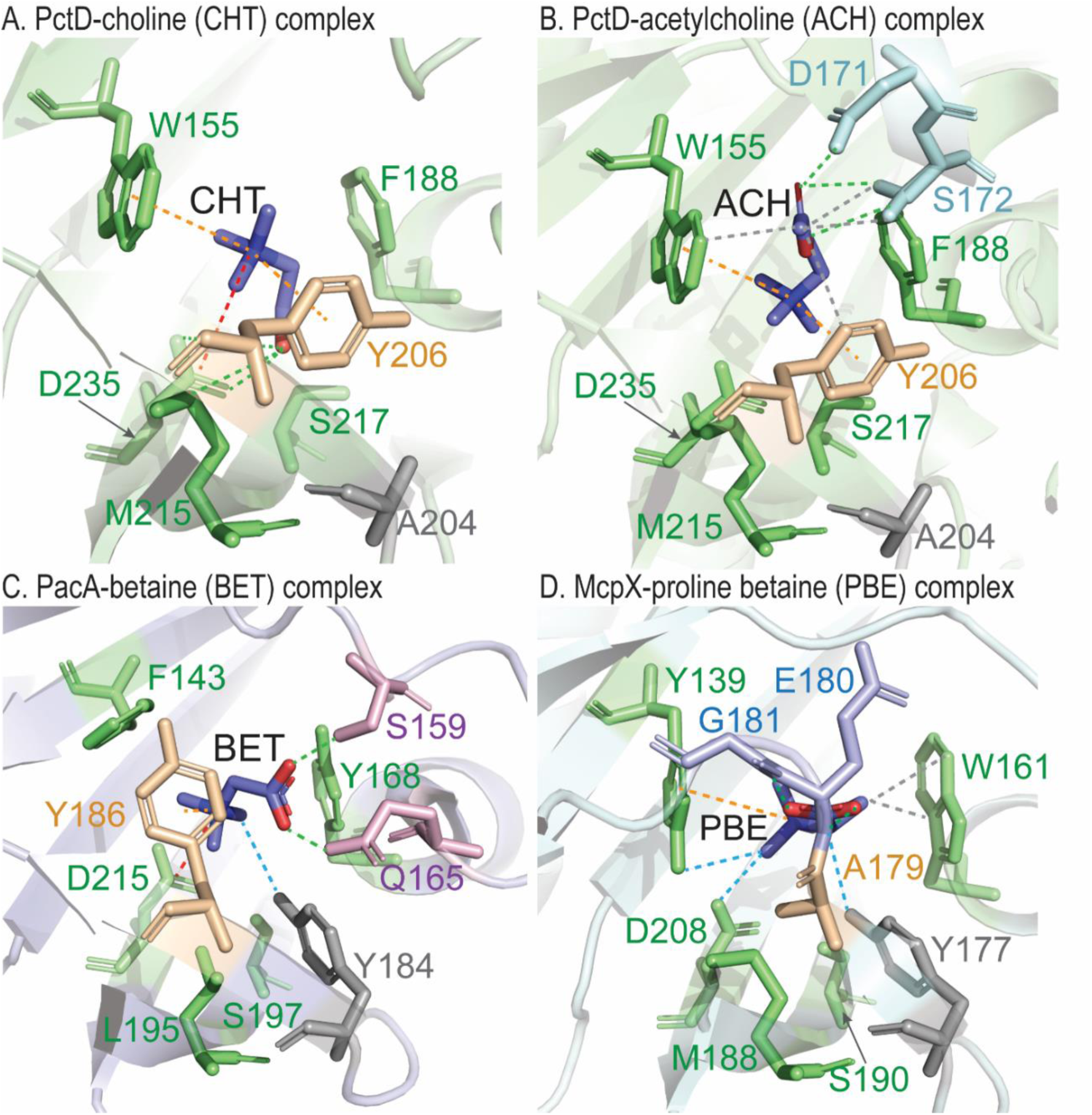
Amine binding receptors have a common ligand-binding interface. **A**. PctD in complex with choline (CHT). **B**. PctD in complex with acetylcholine (ACH). **C**. PacA in complex with betaine (BET). **D**. McpX in complex with proline betaine (PBE). Dashed lines: green – hydrogen bonds, blue – weak hydrogen bonds, orange – cation-π, red – ionic, grey – hydrophobic interactions. Residues shown in green indicate amino acids conserved across structures and analyzed protein sequences. The aromatic residue shown in yellow makes a cation-π bond with ligands. In McpX A179 is in the corresponding position, and it cannot provide such a bond. Another cation-π bond is made with the aromatic residue corresponding to W155 in PctD.

**Figure 2.**
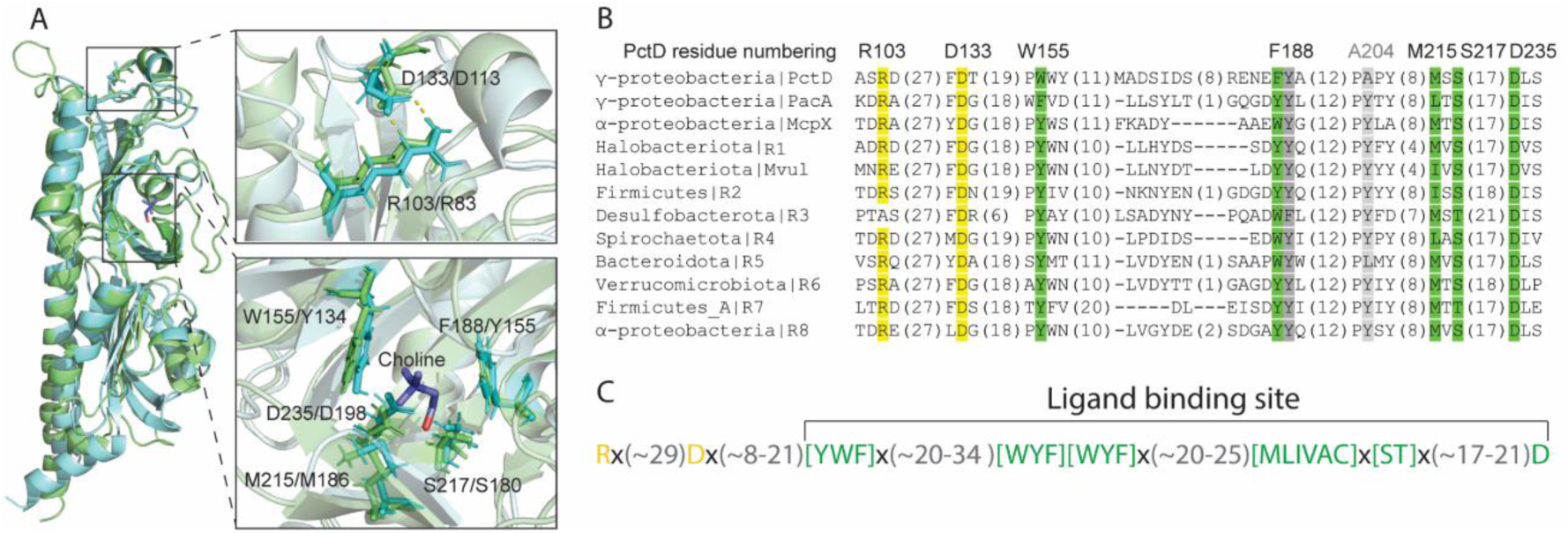
Amine binding sensors have a characteristic signature. **A** Structural superimposition of the PctD sensory domain with the sensory domain of the histidine kinase from the archaeon *Methanosarcina mazei*. PctD is shown in green, the archaeal protein – in cyan. In residue labels in close-up views the first residue corresponds to PctD, the second – to the archaeal protein… **B**. A multiple sequence alignment of amine receptor sensory domain sequences. Residues within the ligand binding site are shown in green and gray, residues in the structure above the ligand binding site are shown in yellow. A residue that is not part of the ligand binding interface, but which exhibits high conservation is shown in dark grey. **C**. The amine binding motif (AM_motif).

### Amine binding dCache_1 domains share a conserved sequence motif.

To find out whether the amine binding site observed in PctD, PacA, and McpX is conserved in other homologous sequences, we performed sensitive profile searches against the NCBI RefSeq database (see Methods) and collected 20,000 dCache_1 domain sequences most similar to PctD, PacA, and McpX. Next, we built a multiple sequence alignment of these sequences and tracked residues forming the amine binding interface in the available structures of quaternary amine-receptor complexes. Through this analysis we identified the most conserved ligand-binding residues, which we tentatively defined as a signature motif for this type of dCache_1 domains (**Fig. 2**). The motif has been identified in the dCache_1 domain of ∼13,000 protein sequences (Dataset S1, multiple sequence alignments are available at https://github.com/ToshkaDev/Amine_motif). Interestingly, the structure of one of these domains, from a histidine kinase of an archaeon *Methanosarcina mazei*, is solved (29), PDB ID 3LIB). We superimposed this structure with that of the quaternary amine binding sensory domain from *P. aeruginosa* PctD and found that overall structures and ligand binding sites are remarkably similar (**Fig. 2A**). Furthermore, residues constituting the motif are located at very similar positions in the bacterial and archaeal proteins (**Fig. 2A**, close-up view, lower panel). We also established that charged residues above the ligand-binding module (R103 and D133 in PctD) are notably conserved (**Fig. 2A**, close-up view, upper panel). Based on these observations we deduced a motif for amine binding receptors (AM_motif) (**Fig. 2C**) and tested it in subsequent experiments.

To verify the contribution of individual conserved amino acid residues of the proposed sequence motif to ligand binding, we prepared alanine substitution mutants of PctD-dCache_1 and submitted them to microcalorimetric titrations with choline using the same experimental conditions that were used for the analysis of the wild type protein (17) (**Fig. 3**). Because significant reduction in the binding affinity was observed for each mutant, experiments were repeated with a higher choline concentration to derive the dissociation constants (Fig. S1). The replacement of the W155 and F188 residues that sandwich the bound ligand resulted in reductions in affinity by factors of 175 and 373, respectively (Table 1). Significant reductions were also observed for Asp235 and Arg103 substitutions (Table 1). Whereas the former residue is part of the binding pocket establishing a hydrogen bond with the bound ligand, Arg103 is outside the binding pocket and likely plays an important role in maintaining the correct geometry of the binding pocket. Replacement of S217 (interaction with choline via a water molecule) and M215 (hydrophobic interaction) had a more modest impact (Table 1). Upon experimental verification, we termed domains that share the motif dCache_1AM.

**Figure 3.**
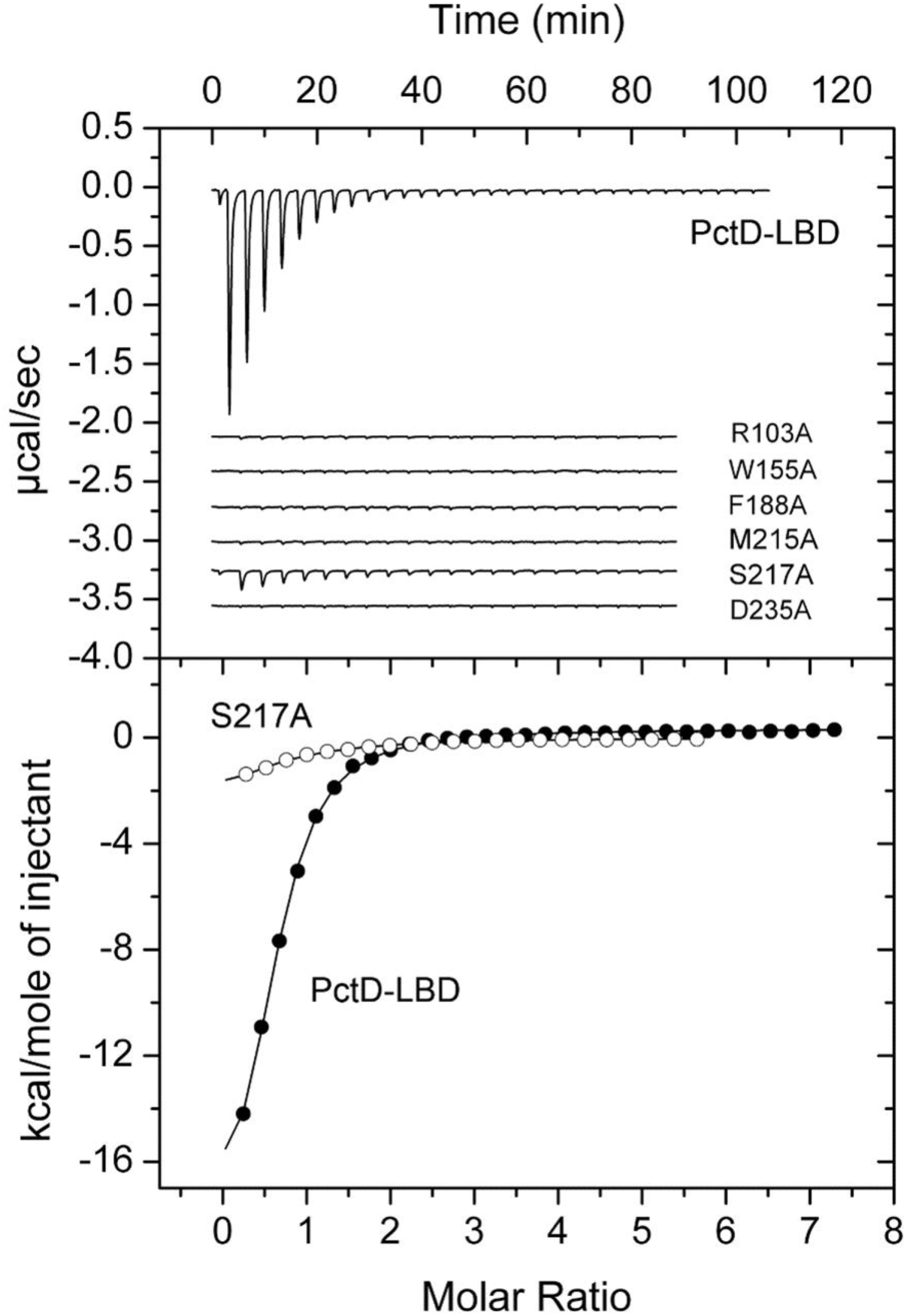
Microcalorimetric titrations of the PctD sensor domain and site directed mutants in individual residues of the amine binding motif with choline. Upper panel: Raw data for the titration of 16 µM of protein with 9.6 µl aliquots of 0.5 µM choline. Lower panel: Concentration-normalized and dilution heat corrected integrated raw data. The line is the best fit using the “One binding site model” of the MicroCal version of ORIGIN. In case no binding heats were observed experiments were repeated with a higher ligand concentration. The resulting curves are shown in Fig. S1 and the derived dissociation constants are provided in Table 1.

**Table 1).** Microcalorimetric studies for the binding of choline to PctD-LBD and site-directed mutants in amino acids of the QA binding motif. The corresponding titration curves are shown in Fig. S1.

| Protein | $K_D$ (μM) | Reduction with respect to wt protein (fold) |
| --- | --- | --- |
| PctD-LBD<sup>a</sup> | $2.6 \pm 0.1$ | - |
| PctD-LBD_D235A | $421 \pm 17$ | 162 |
| PctD-LBD_F188A | $970 \pm 117$ | 373 |
| PctD-LBD_W155A | $454 \pm 20$ | 175 |
| PctD-LBD_M215A | $150 \pm 17$ | 58 |
| PctD-LBD_S217A | $14.2 \pm 1.4$ | 5.4 |
| PctD-LBD_R103A | $943 \pm 54$ | 363 |

<sup>a</sup> reported previously in (17)

### dCache_1AM domains are widespread and found in all major receptor types

We identified dCache_1AM domains in more than 13,000 proteins from 8,000 species of bacteria and archaea (Datasets S1 and S2). We have extracted the corresponding isolation sources from the NCBI BioSample section of NCBI (Dataset S2) and observed that many dCache_1AM domains come from plant-associated bacteria, both beneficial, such as *Sinorhizobium meliloti* and *Azospirillum brasilense*, and pathogenic, such as *Xanthomonas campestris*. Similarly, we identified dCache_1AM domains in the human microbiome, including both beneficial human gut bacteria, for example *Roseburia intestinalis* and *Ruminococcus lactaris*, and human pathogens, such as *Campylobacter jejuni* and *Aeromonas hydrophila* (**Fig. 4**, Dataset S2). Analysis of dCache_1AM phyletic distribution revealed that receptors containing this sensor domain are found in many bacterial and in one archaeal phylum – Halobacteriota (**Fig. 4A**, Dataset S2).

**Figure 4.**
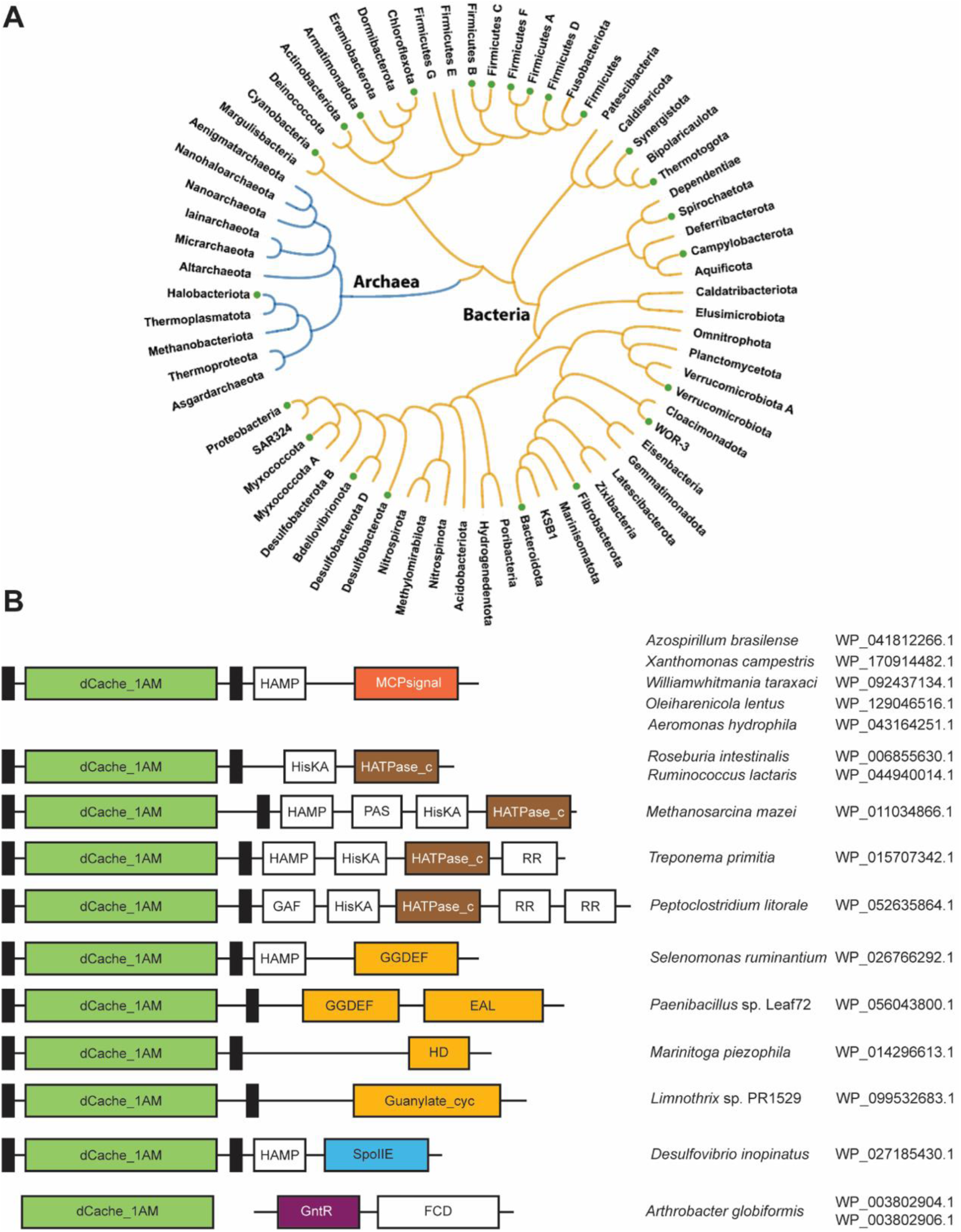
Phyletic distribution (**A**) and prevalent domain architectures of amine receptors (**B**). Solid circles at the tips of the tree branches indicate that the AM_motif was found in the corresponding phylum. Domain definitions follow the Pfam domain nomenclature (InterPro ids are in parentheses): EAL (IPR035919), a diguanylate phosphodiesterase; GGDEF (IPR000160), a diguanylate cyclase; Guanylate_cyc (IPR001054), an adenylate or guanylate cyclase; HATPase_c (IPR003594), a histidine kinase; HD (IPR006674), phosphohydrolase; MCPsignal (IPR004089), methyl-accepting chemotaxis protein (chemoreceptor); SpoIIE (IPR001932), serine/threonine phosphatase; GntR (IPR000524), transcription regulator.

By performing domain analysis of all dCache_1AM-containing proteins we revealed that they are exclusively found in signal transduction proteins or as stand-alone domains (**Fig. 4B**, Dataset S1). We found dCache_1AM domains in all four major types of bacterial and archaeal transmembrane receptors: chemoreceptors, sensor histidine kinases, cyclic (di)-nucleotide turnover enzymes and serine/threonine phosphatases (**Fig. 4B**, Dataset S1). In bacteria, the vast majority of dCache_1AM domains are found in chemoreceptors, whereas in archaea they are predominantly found in sensor histidine kinases. We also identified an unusual case of an intracellular dCache_1AM domain in Actinobacteria. In many actinobacterial genomes, including representatives of *Streptomyces*, *Nocardia*, *Rhodococcus*, *Arthrobacter, Mycobacterium* and other genera, a stand-alone dCache_1AM domain protein is encoded in a two-gene operon with a transcription factor (**Fig. 4B**, Dataset S2). This is a rare example of repurposing an extracellular domain for intracellular sensing.

### dCache_1AM domains bind primary, secondary, tertiary, and quaternary amines

From the list of more than 13,000 dCache_1AM sequences (Dataset S1), we selected ten targets (R1 through R10) for experimental verification (Table 2). These domains were selected from (i) four major receptor families, namely sensor histidine kinases, chemoreceptors, Ser/Thr phosphatases, and diguanylate cyclases/phosphodiesterases and (ii) one archaeal and several bacterial phyla (Table 2). Two targets, each containing a single amino acid substitution in the AM_motif, were used as a negative control (R9 and R10, Table 2). The individual dCache_1AM domains from these ten receptors were overexpressed in *Escherichia coli* and purified by affinity chromatography. For nine of these proteins a buffer system was identified that guaranteed protein solubility and stability, whereas the remaining protein, R2, was insoluble despite several solvent engineering efforts. Thermal unfolding studies revealed a transition for all other proteins, which is indicative of protein folding. Because the AM_motif was established based on the binding of quaternary amines, thermal shift unfolding studies and isothermal titration experiments were conducted in parallel to study the binding of various quaternary amines. We have conducted ITC experiments for ligands that caused an increase in the midpoint of proteins unfolding (Tm) by at least 2 °C. We were able to observe the binding of choline and/or acetylcholine to four of the eight target proteins and one target protein, R5, bound trimethylamine N-oxide (TMAO) (**Fig. 5**, Fig. S2, Table 3).

**Table 2).**
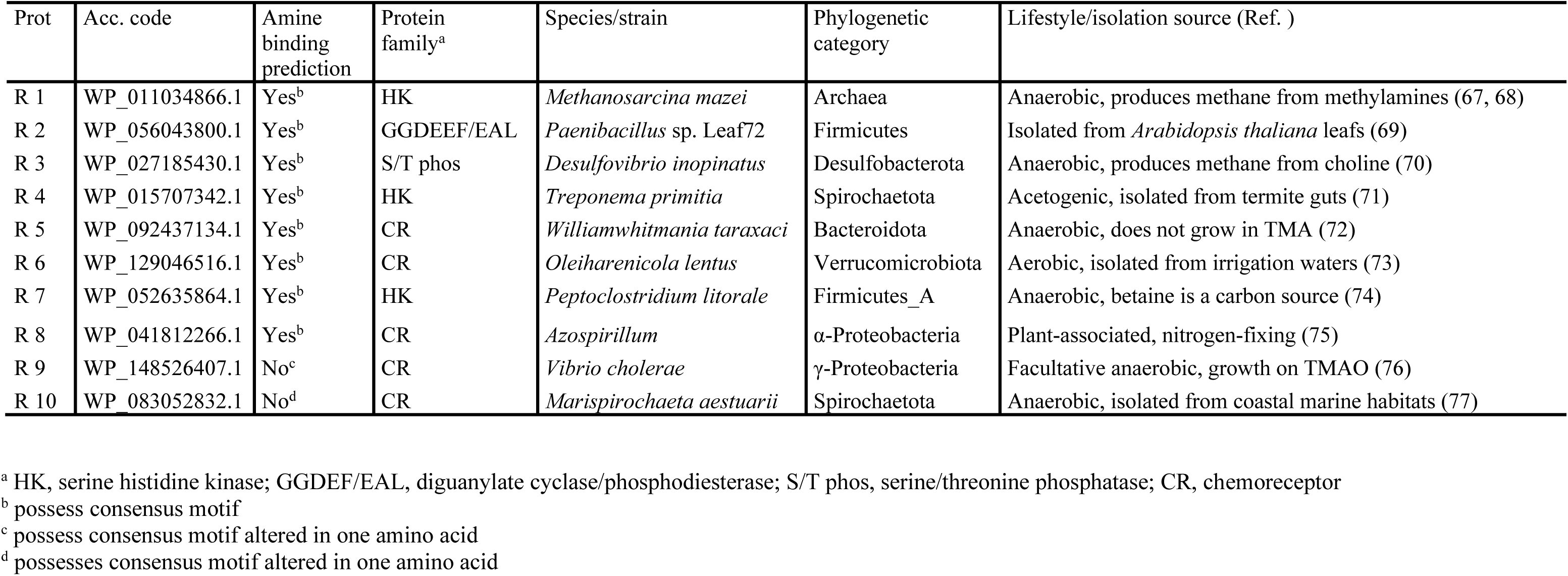
Characteristics of the 10 receptor proteins analyzed in this study. Proteins R 1 to R 8 contain the amine binding motif and are predicted to bind amines. Proteins R 9 and R 10 possess slight modification of this binding motif (specified beneath the Table) and were predicted to not bind amines.

| Prot | Acc. code | Amine binding prediction | Protein family<sup>a</sup> | Species/strain | Phylogenetic category | Lifestyle/isolation source (Ref. ) |
| --- | --- | --- | --- | --- | --- | --- |
| R 1 | WP_011034866.1 | Yes<sup>b</sup> | HK | *Methanosarcina mazei* | Archaea | Anaerobic, produces methane from methylamines (67, 68) |
| R 2 | WP_056043800.1 | Yes<sup>b</sup> | GGDEEF/EAL | *Paenibacillus* sp. Leaf72 | Firmicutes | Isolated from *Arabidopsis thaliana* leafs (69) |
| R 3 | WP_027185430.1 | Yes<sup>b</sup> | S/T phos | *Desulfovibrio inopinatus* | Desulfobacterota | Anaerobic, produces methane from choline (70) |
| R 4 | WP_015707342.1 | Yes<sup>b</sup> | HK | *Treponema primitia* | Spirochaetota | Acetogenic, isolated from termite guts (71) |
| R 5 | WP_092437134.1 | Yes<sup>b</sup> | CR | *Williamwhitmania taraxaci* | Bacteroidota | Anaerobic, does not grow in TMA (72) |
| R 6 | WP_129046516.1 | Yes<sup>b</sup> | CR | *Oleiharenicola lentus* | Verrucomicrobiota | Aerobic, isolated from irrigation waters (73) |
| R 7 | WP_052635864.1 | Yes<sup>b</sup> | HK | *Peptoclostridium litorale* | Firmicutes_A | Anaerobic, betaine is a carbon source (74) |
| R 8 | WP_041812266.1 | Yes<sup>b</sup> | CR | *Azospirillum* | α-Proteobacteria | Plant-associated, nitrogen-fixing (75) |
| R 9 | WP_148526407.1 | No<sup>c</sup> | CR | *Vibrio cholerae* | γ-Proteobacteria | Facultative anaerobic, growth on TMAO (76) |
| R 10 | WP_083052832.1 | No<sup>d</sup> | CR | *Marispirochaeta aestuarii* | Spirochaetota | Anaerobic, isolated from coastal marine habitats (77) |

<sup>a</sup> HK, serine histidine kinase; GGDEF/EAL, diguanylate cyclase/phosphodiesterase; S/T phos, serine/threonine phosphatase; CR, chemoreceptor
<sup>b</sup> possess consensus motif
<sup>c</sup> possess consensus motif altered in one amino acid
<sup>d</sup> possesses consensus motif altered in one amino acid

**Figure 5.**
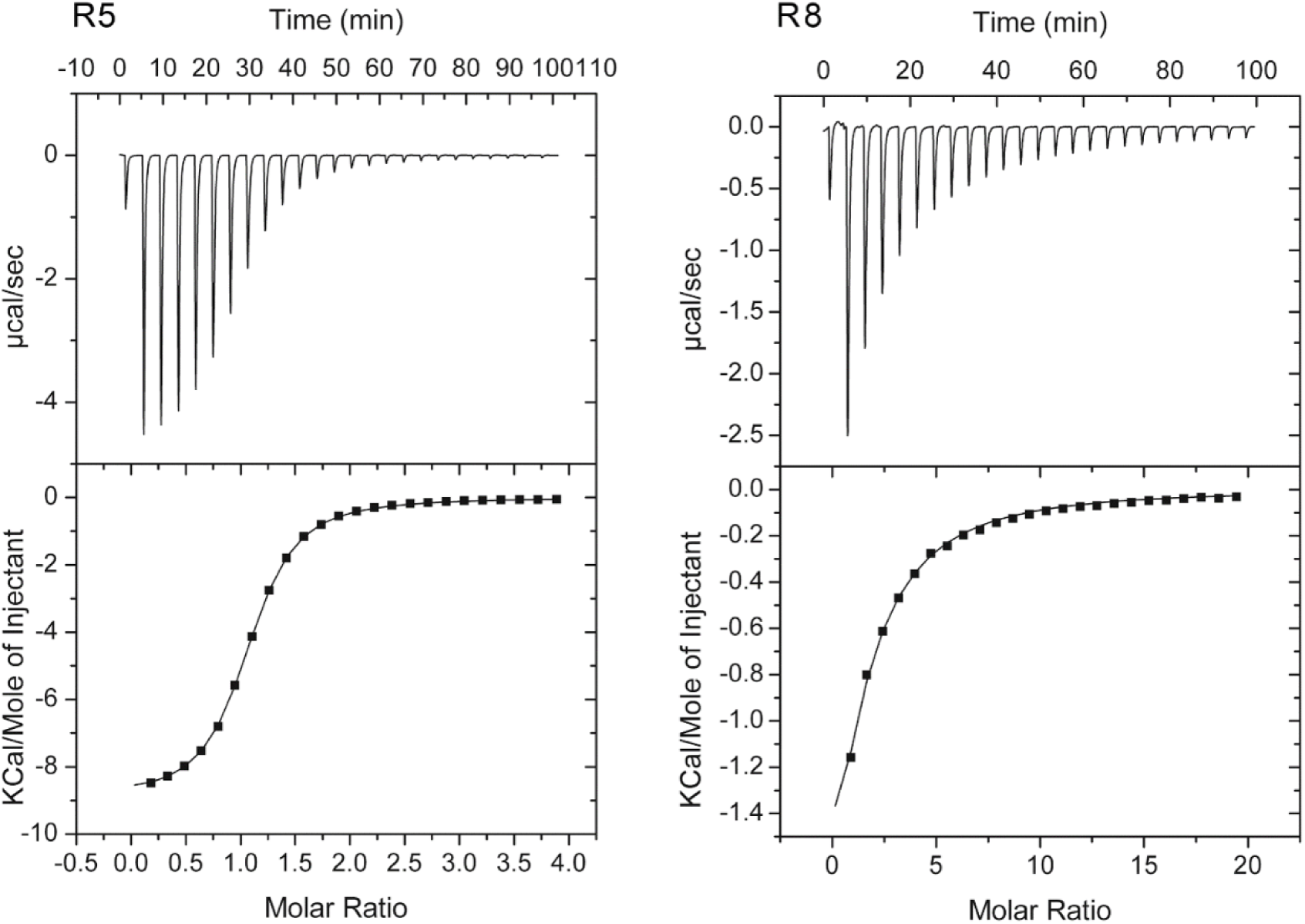
Microcalorimetric titrations of predicted amine responsive dCache domains with choline. Upper panel: Raw data for the titration of 75 µM of protein with 8.0 µ1 aliquots of 2 mM (R5) or 10 mM (R8) choline. Lower panel: Concentration-normalized and dilution heat corrected integrated raw data. The line is the best fit using the “One binding site model” of the MicroCal version of ORIGIN.

**Table 3).** Dissociation constants derived from microcalorimetric binding studies of different amines to sensor domains of the 10 receptors analyzed. **Dissociation constants below 10 µM are shown in bold face.**

| Protein | $K_D$ (μM) | | | | | | | | |
| --- | --- | --- | --- | --- | --- | --- | --- | --- | --- |
|  | Quaternary amines |  |  | Methyl-/ethyl-/ethanol amines<sup>a</sup> |  |  |  |  | Polyamine |
|  | Acetylcholine | Choline | TMAO | Methylamine | Ethylamine | Dimethylamine | Trimethylamine | Ethanolamine | Ethylenediamine |
| R1<sup>b</sup> | 1845 ± 1120 | Nb<sup>c</sup> | Nb | 35.1 ± 3 | 19.6 ± 2 | **0.53 ± 0.02** | 10.0 ± 0.4 | Nb | Nb |
| R2 | Protein insoluble |  |  |  |  |  |  |  |  |
| R3<sup>d</sup> | Nb | Nb | Nb | Nb | Nb | Nb | Nb | **2.3 ± 0.1** | 29.6 ± 3 |
| R4<sup>e</sup> | Nb | Nb | Nb | Nb | Nb | Nb | Nb | Nb | Nb |
| R5<sup>b</sup> | 581 ± 129 | **3.6 ± 0.1** | 1742 ± 88 | Nb | Nb | 311 ± 79 | **1.2 ± 0.1** | Nb | Nb |
| R6<sup>b</sup> | Nb | Nb | Nb | Nb | Nb | Nb | 151 ± 4 | Nb | Nb |
| R7<sup>d</sup> | Nb | 2950 ± 217 | Nb | 16.6 ± 2 | Nb | **5.3 ± 0.4** | 12.6 ± 1 | **1.7 ± 0.02** | 12.3 ± 1 |
| R8<sup>b</sup> | 735 ± 24 | 183 ± 4 | Nb | 25.1 ± 3 | 46.1 ± 7 | **5.8 ± 1** | 24.0 ± 4 | Nb | Nb |
| R9<sup>d</sup> | Nb | Nb | Nb | Nb | Nb | Nb | Nb | Nb | Nb |
| R10<sup>d</sup> | Nb | Nb | Nb | Nb | Nb | Nb | Nb | Nb | Nb |
| PacA-LBD | Nb<sup>f</sup> | 113 ± 16<sup>f</sup> | Nd<sup>g</sup> | Nb | Nb | Nb | 113 ± 6 | Nb | Nb |

<sup>a</sup> Volatile organic compound
<sup>b</sup> No binding by ITC of L-Carnitine, betaine, proline and trigonelline
<sup>c</sup> No binding by ITC and/or Thermal shift assay (increases above 2°C were considered significant).
<sup>d</sup> No binding by ITC of L-Carnitine and trigonelline
<sup>e</sup> No binding by ITC of L-Carnitine, betaine and trigonelline
<sup>f</sup> Reported previously in (17)
<sup>g</sup> Not determined

Surprisingly, none of the other quaternary amines caused any significant increases in Tm. Since thermal unfolding experiments at times give false-negative results (i.e. ligand binding that does not significantly increase the Tm) we conducted isothermal titration experiments with the maximal possible concentration (10 to 20 mM) of other biologically important quaternary amines, such as L-carnitine, betaine, proline and trigonelline (Footnotes to Table 3). However, we observed no binding in any of these experiments, suggesting that choline and acetylcholine are the main quaternary amine ligands recognized by dCache_1AM-containing receptors.

Four of the selected targets failed to bind quaternary amines and in subsequent studies we aimed at establishing their ligands. Target R1 came from an archaeon *Methanosarcina mazei*, which uses methylamines as carbon and energy sources (30). We used the available structure of R1 (PDB ID: 3lib) (29) to conduct *in silico* docking experiments with methylamine, dimethylamine, and trimethylamine and found that all ligands bind to the same ligand binding pocket of the target; we observed that methylamine and dimethylamine make contacts with the conserved residues constituting the AM_motif (Fig. S3). Subsequently, we have conducted ligand screening with the target proteins and various small biogenic amines. We observed significant TM increases for methylamine, ethylamine, ethylenediamine, dimethylamine, trimethylamine and ethanolamine using the thermal shift assay (Table S1). Six out of the seven target proteins showed a significant increase in Tm in the presence of small biogenic amines. We subsequently conducted ITC experiments to derive the corresponding dissociation constants (**Fig. 6**, Fig. S4). Three proteins, R1, R7 and R8, showed a wide ligand spectrum and recognized with high affinity four of the amine compounds. Proteins R1 and R8, that come from an archaeon and an alphaproteobacterium, had the same ligand profile. Both proteins had a preference for dimethylamine but also recognized with lower affinity methyl-, ethyl- and trimethylamine (Table 3). Other proteins showed a narrower ligand spectrum; for example, R5 recognized trimethylamine with high affinity, but its affinity for dimethylamine was reduced ∼300-fold (Table 3). In addition, R5 bound choline with an affinity very similar to that of trimethylamine, indicative of plasticity in ligand recognition (Table 3).

**Figure 6.**
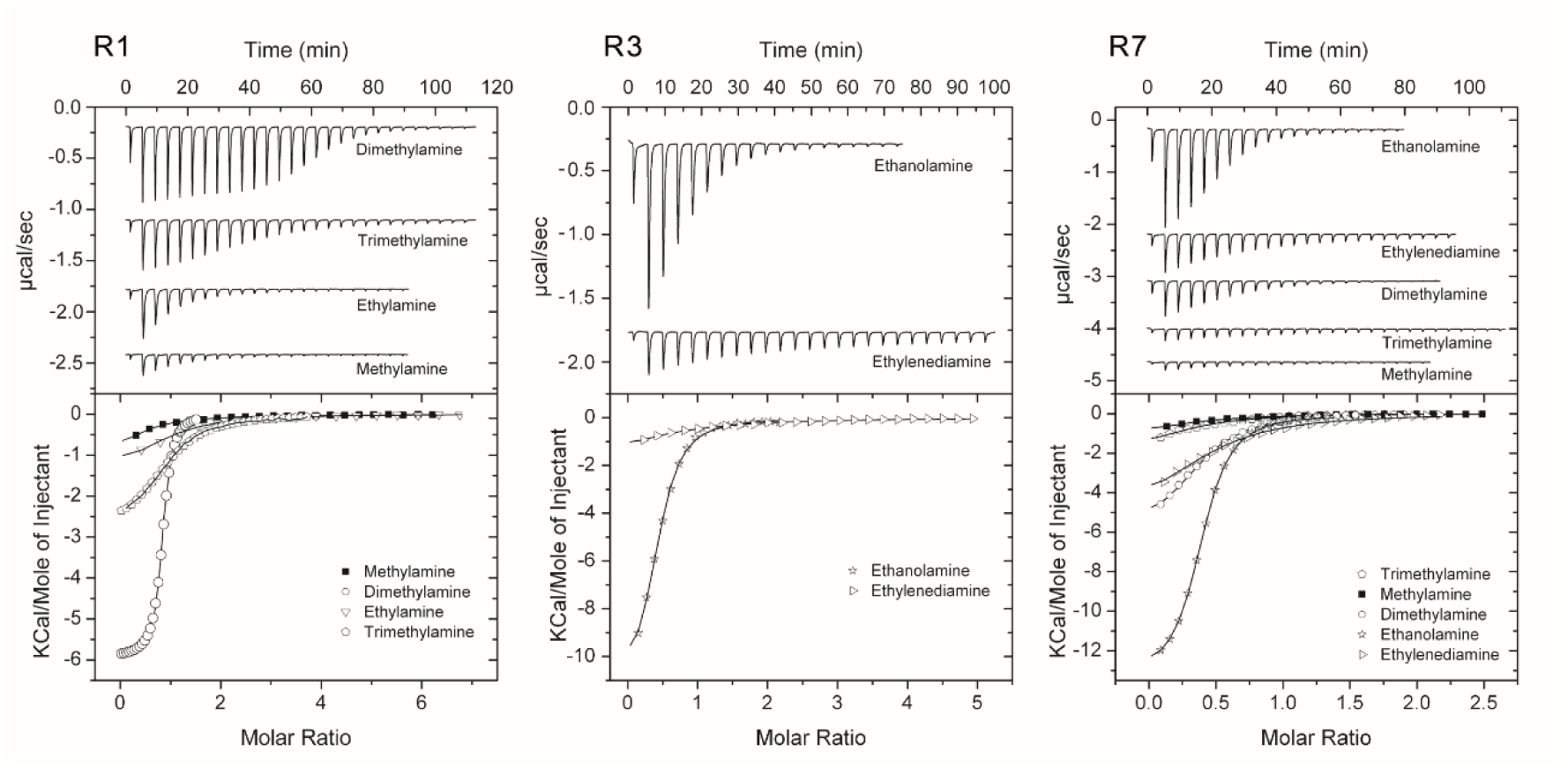
Microcalorimetric titrations of predicted amine responsive dCache domains with small biogenic amines. Upper panel: Raw data for the titration of 30 to 50 µM of protein with 3.2 to 11.1 µl aliquots of 1 to 2 mM amine solutions. Lower panel: Concentration-normalized and dilution heat corrected integrated raw data. The line is the best fit using the “One binding site model” of the MicroCal version of ORIGIN.

The only predicted amine binding dCache_1 domain for which no binding was observed in our experiments was R4. However, it cannot be ruled out that this protein binds other amine(s) that we have not tested. As mentioned above, targets R9 and R10 were used as a negative control: both contained a single mismatch in the AM_motif. Thermal shift assays and ITC experiments with the maximal possible ligand concentration did not provide evidence for the binding of any of these ligands, further validating the conserved AM-motif. Based on these results, we wanted to verify whether the quaternary amine sensing chemoreceptor PacA (17) also binds methylamines that were not tested in the previous study (17). Thermal shift (Table S1) and microcalorimetric titrations (Table 3) revealed that indeed PacA also binds trimethylamine, further suggesting that the capacity to bind various biogenic amines – primary, secondary, tertiary, and quaternary - is a general property of the dCache_1AM domain family.

### dCache_1AM domains evolved from the universal amino acid sensor

To establish the evolutionary origins of the dCache_1AM domains we performed sequence, structure, and phylogenetic analyses. Analysis of the multiple sequence alignment showed that AM- and AA motifs have several overlapping positions (**Fig. 7A**, the alignment in FASTA format is available at https://github.com/ToshkaDev/Amine_motif). Two aromatic residues (corresponding to F188 and Y189 in PctD) in the middle of the motifs and aspartate at the end of the motifs (corresponding to D235 in PctD; see **Fig. 7A**) are conserved in both motifs. Another position highly conserved in amino acid receptors as aromatic position is shared with amine receptors (corresponds to A204 in PctD), although the position is more variable in dCache_1AM. In contrast, an aromatic and a positively charged positions at the beginning of the AA motif are not conserved in the AM motif. dCache_1AM domains, in addition, have an insertion upstream the ligand binding site, which includes charged residues R103 and D133 that make hydrogen bonds with each other (see **Fig. 7A**, **2A**). Two positions of the AM motif (corresponding to M215 and S217 in PctD) are also found in amino acid receptors, although while in dCache_1AM the M215 position is exclusively hydrophobic, in dCache_1AA this position is more variable (**Fig. 7A**, **Fig. 2B**).

**Figure 7.**
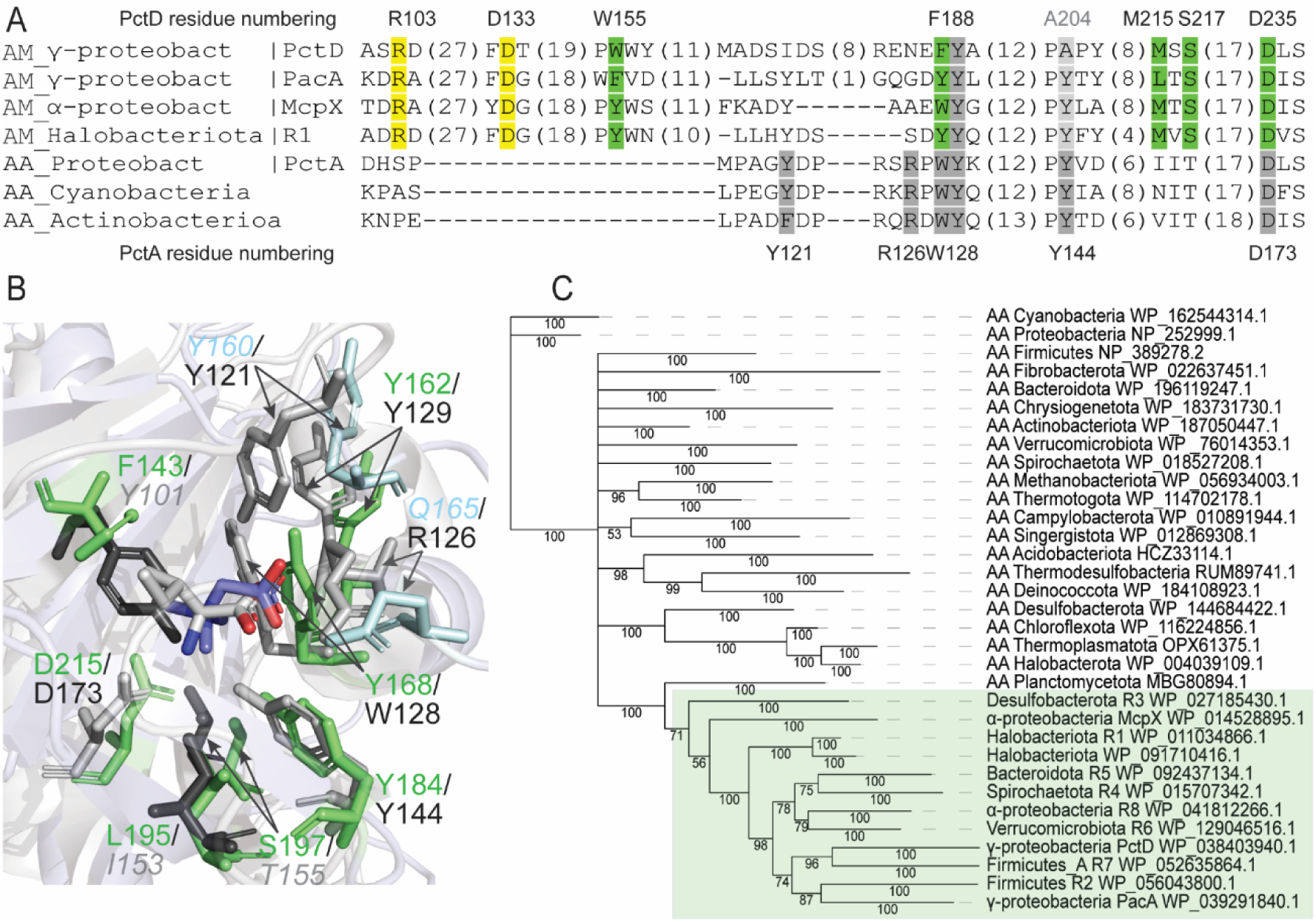
Amine receptors evolved from amino acid receptors. **A.** Multiple sequence alignment of amine and amino acid receptor sensory domains. **B**. Structural superimposition of ligand binding pockets of the amine receptor PacA from *P. atrosepticum* and the amino acid receptors PctA from *Pseudomonas aeruginosa*. Residues corresponding to the AM_motif are shown in green, residues in grey correspond to the AA_motif. Residues in the PacA pocket making hydrogen bonds with betaine are shown in blue. Residues that are not part of the corresponding motif but that are in positions equivalent to the residues in the other motif are shown with a faint italic font. **C**. Bayesian phylogenetic tree of amine and amino acid receptor sensory domains. Amine receptors are shown in green background.

To explore structural similarities, we superimposed ligand binding pockets of dCache_1AA and dCache_1AM and observed similar ligand binding interfaces (**Fig. 7B**). Key residues constituting both the AM and AA motifs are in the same positions. One of the key residues of the AA_motif, corresponding to D173 in PctD, is well conserved in amine receptors; quaternary amines in PctD-CHT, PacA-BET, McpX-PBE complexes, as well as docked methylamines in the archaeal protein make contacts with this residue (see **Fig. 1**, **Fig. 7AB**, and Fig. S3). Two shared aromatic residues (Y168/W128 and Y184/Y144) are in favorable positions for ligand binding in both amine and amino acid receptors. A structurally important and highly conserved aromatic residue, which does not interact with the ligand (Y162/Y129) is found in a similar position in both amine-binding PacA and amino acid-binding PctA (**Fig. 7B**). The main difference between AA and AM motifs was observed in three positions. A key position of the AA_motif corresponding to R126 in PctA makes a hydrogen bond with the carboxyl group of amino acid ligands. This position is not conserved in amine receptors and in its closest structural equivalent in PacA is Q165, which also makes a hydrogen bond with its designated ligand, betaine (**Figs. 7B and 1C**). An aromatic residue Y160 corresponding to Y121 in PctA is oriented in the opposite direction in PacA and does not contribute to ligand coordination (**Fig. 7B**). Conversely, an important aromatic residue making a cation-π bond with quaternary amine ligands is oriented downward in amino acid receptors and cannot interact with amino acid ligands (**Fig. 7B**).

We then inferred a Bayesian phylogenetic tree using protein sequences of dCache_1AA and dCache_1AM from several bacterial and archaeal phyla (see Methods, the tree in NEXUS format is available at https://github.com/ToshkaDev/Amine_motif). The tree showed that all dCache_1AM sequences are found in a single branch derived from one of the branches of a more diverse set of dCache_1AA sequences (**Fig. 7C**). An amino acid receptor from Planctomycetota was the closest to amine binding sensors among the ones used for the phylogenetic inference.

## Discussion

Prior to this work, three bacterial chemoreceptors from closely related proteobacterial species were shown to bind and respond to quaternary amines and polyamines as signaling molecules (15–18). In this study, we identify thousands of bacterial and archaeal receptors containing dCache_1AM domains that bind various biogenic amines. We computed and experimentally verified a conserved sequence motif signature for this novel class of sensory domains. We show that dCache_1AM sensors bind not only quaternary, but also primary, secondary, and tertiary amines as well as the polyamine ethylenediamine. Furthermore, based on their distribution and superior affinity, we conclude that small biogenic amines rather than quaternary amines constitute the primary ligand group for the dCache_1AM class (Table 3).

We show that amine sensors evolved from the universal dCache_1AA amino acid sensors (8) through a small insertion in the ligand binding pocket and replacement of key ligand binding residues. Although amine sensors are not as ubiquitous as amino acid sensors, they are still widespread. These domains are identified in all major types of bacterial transmembrane receptors - chemoreceptors, sensor histidine kinases, serine/threonine phosphatases and cyclic (di)nucleotide turnover enzymes – from several major bacterial and one archaeal phyla. In archaea we identified them in two classes of Halobacteriota -Methanomicrobia and Methanosarcinia - where they were horizontally transferred from bacteria. Structural analyses of bacterial and archaeal receptors showed that the ligand binding interface is well conserved in phylogenetically distant species.

Amine compounds sensed by dCache_1AM receptors are of enormous ecological importance. Many biogenic amines serve as energy, carbon and nitrogen sources for bacteria (31, 32). Quaternary amines, such as acetylcholine, are important neurotransmitters and mediators of inter-kingdom and inter-bacterial interactions (33). Methyl-/ethyl-/ethanol amines are volatile organic compounds (voc). Voc are known to mediate plant-bacteria interactions and bacteria-derived voc were found to promote the growth, health, immunity and stress resistance of plants (34–37). However, the molecular detail of voc signaling is frequently lacking. The presence of dCache_1AM domains in many plant-associated bacteria (Dataset S2), suggests a potential role of the corresponding receptors in mediating plant-bacteria interactions.

The fact that the domain family identified responds to QAs as well as to methyl-/ethyl-/ethanol amines is the consequence of i) an important structural similarity of these ligands (Fig. S5A) and ii) an intertwining of their metabolism (Fig. S5B). For example, methanogenic archaea use methylamine, dimethylamine, trimethylamine, betaine and choline as carbon and energy sources (38). Methylamine and trimethylamine are the result of betaine (24) and choline degradation (39), respectively. In humans, the production of TMAO from trimethylamine is linked to multiple diseases such as trimethylaminuria, atherosclerosis or cardiovascular disease (40, 41). In marine environments, trimethylamine is produced from TMAO, glycine betaine, choline and carnitine (42, 43). Dimethylamine and trimethylamine are produced in sewage water from choline and creatinine, which are present in urine as well as plant and animal tissues (44).

Of all tested ligands and target sensor domains, the highest binding affinity was observed for the binding of dimethylamine to dCache_1AM domain (target R1) that is found in a histidine kinase from *Methanosarcina mazei* (Table 3). Interestingly, R1 was among the first reported 3D structures of the Cache superfamily (29); however, ligands recognized by this domain remained unknown until now*. M. mazei* is a model methanogenic archaeon, found primarily in sewages and anoxygenic environments that are sources of quaternary and methylated amines (24). *M. mazei* has multiple genes encoding enzymes that demethylate trimethylamine (*mttB*), dimethylamine (*mtbB*) and methylamine (*mtmB*) (45), which is the initial event of a multi-step process leading to methane generation of. We found that R1 binds all three substrates of the demethylation reactions with high affinity (Table 3). The fact that R1 also binds ethylamine (Table 3) suggests that this compound may also serve as a substrate for the demethylation reactions. The R1-containing sensor histidine kinase is encoded in a gene cluster with a dimethylamine methyltransferase (Fig. S6) and, coincidently, dimethylamine is a preferred ligand for R1 (Table 3). Transcript levels of this methyltransferase are significantly upregulated in the presence of trimethylamine (45), which is the second best substrate for R1, and it is likely that the R1-containing histidine kinase is involved in this regulatory process.

The successful prediction of ligands for sensor domains that recognized their ligands through different types of bonding interactions, i.e. primarily H-bonds or hydrophobic interactions, indicates that this procedure appears to be generally applicable to identify ligands recognized by sensor domains. These approaches will allow the annotation of thousands of receptor proteins with a cognate ligand class. Such information will have a major impact in the field since it will enable precise wet-lab experiments to define the function of a given regulatory system. Thus, our approach holds the strong promise to address an important bottleneck in microbiology (5): identification of signal molecules that stimulate numerous signal transduction cascades in bacteria and archaea.

## Materials and Methods

### Strains and plasmids

The bacterial strains and plasmids used are listed in Table S2.

### Identification of the amine motif containing dCache domains

Amine binding protein sequences were identified in the following two steps. The NCBI RefSeq protein database was downloaded (July 2022) and searched with the dCache_1 domain profile hidden Markov model (the InterPro (46) identifier IPR033479) with the E-value threshold of 0.01 both for sequences and domains. Protein sequence regions corresponding to the dCache_1 domain were extracted from the identified sequences and divided into four separate datasets, and each was aligned on the local computational cluster using the FFT-NS-2 algorithm of the MAFFT package (47). Then, in each dataset, the positions corresponding to the defined AM_motif were tracked and corresponding protein sequences were extracted. In parallel, PSI-BLAST searches were initiated against the NCBI RefSeq database using the McpX, PacA, and PctD dCache_1 domain sequences with the maximal number of sequences set to 20,000. The obtained protein dCache_1 domain sequences were merged and aligned using the FFT-NS-2 algorithm of the MAFFT package. In the aligned protein set, positions corresponding to the AM_motif were tracked and corresponding protein sequences were extracted. At the final step, protein sequences extracted from the results of the HMM and PSI-BLAST searches were combined and realigned using the L-INS-i algorithm of the MAFFT package and again the AM_motif positions were tracked and verified. The GTDB taxonomy for the final sequence set was retrieved using the GTDB metadata tables (https://data.ace.uq.edu.au/public/gtdb/data/releases/release202/202.0/).

### Multiple sequence alignment, domain identification

Jalview (48) was used to explore and edit the alignments. Domains were identified running TREND (49, 50) with the Pfam profile HMMs. The generated data were downloaded in JSON format from the website and processed programmatically to determine domain architecture variants and abundances. Additional sensitive profile-profile searches were carried out using HHpred (51).

### Phylogeny inference

The sequence alignment was edited using an alignment trimming tool, trimAl (52): positions in the alignment with gaps in 10% or more of the sequences were removed unless this leaves less than 60%. In such case, the 60% best (with fewer gaps) positions were preserved. The amino acid replacement model for the set of protein sequences was determined running ProtTest (53) and based on Akaike (54) and Bayesian (55) information criteria. The best model was found to be LG with gamma distribution of rate variation across sites in combination with the proportion invariable site model and empirical state frequencies (LG + I + G + F). LG is an improved model compared to WAG, which has been achieved by incorporating the variability of evolutionary rates across sites in the matrix estimation and using a much larger and diverse database (56). Using the determined amino acid replacement model, a phylogenetic tree was inferred using a bayesian inference algorithm implemented in MrBayes (57). Metropolis-coupled Markov chain Monte Carlo simulation implemented in MrBayes was run with 3 heated and 1 cold chain and discarding the first 25% of samples from the cold chain at the “burn-in” phase. 900,000 generations were run till the sufficient convergence was achieved (the average standard deviation of split frequencies equal to 0.01) with chain sampling every 500 generations.

### Protein structure manipulations

Protein structures of the target proteins were modeled using AlphaFold 2 (58). Comparative analysis of solved and modeled protein structures was done using PyMoL (59) and Mol* Viewer (60).

### In silico docking

AutoDock Vina (61) was used for computational docking experiments. The protein structure of the histidine kinase dCache_1 domain from the archaeon *Methanosarcina mazei* (PDB ID 3LIB) was prepared using MGLTools. For the experiments, we downloaded ligands from the Zink database (62) in mol2 format and prepared them for the analysis using the Open Babel toolbox (63) and custom shell script. The docking was performed with the search exhaustiveness 8. Coordinates of the center of the simulation box (Angstroms): X: -25.388; Y: -45.395; Z: -2.575, b) the box dimensions (Angstroms): X: 20; Y: 24; Z: 20.

### Protein overexpression and purification

The transmembrane regions of proteins R1 to R10 were determined using TMHMM (64). pET28b(+) expression plasmids encoding the region between both transmembrane regions (i.e. the LBD) fused to an N-terminal His-tag were purchased from GeneScript Biotech (Netherlands). The corresponding protein sequences are provided in Table S3. The site-directed mutants of PctD-LBD were purified like the native protein (17). The remaining proteins were purified as published previously (65) using the buffers specified in Table S4. Freshly purified proteins were dialyzed overnight into the buffers specified in Table S4 for immediate analysis.

### Thermal shift assays

The detailed experimental protocol of the thermal shift assays has been reported in (66). Briefly, assays were carried out using a MyIQ2 Real-Time PCR instrument (BioRad, Hercules, CA, USA). Experiments were conducted in 96-well plates and each assay mixture contained 20.5 μL of the dialyzed protein (10 – 70 µM), 2 μL of 5 X SYPRO orange (Life Technologies, Eugene, Oregon, USA) and 2.5 μL of the 20 mM ligand solution or the equivalent amount of buffer in the ligand-free control. Samples were heated from 23 °C to 85°C at a scan rate of 1 °C/min. The protein unfolding curves were monitored by detecting changes in SYPRO Orange fluorescence. The Tm values correspond to the minima of the first derivatives of the raw fluorescence data.

### Isothermal Titration Calorimetry (ITC)

Experiments were conducted on a VP-microcalorimeter (Microcal, Amherst, MA). The site-directed mutants of PctD-LBD were analyzed as reported previously (17). In the case no binding heats were observed, experiments were repeated with a higher choline concentration and the resulting curves (Fig. S1) were used to derive the dissociation constants (Table 1). Proteins R1 to R10 were dialyzed into the analysis buffer, placed at a concentration of 10 to 75 µM into the sample cell and titrated with freshly made up amine solutions (0.5 to 20 mM). In the case no binding heats were observed for a titration with 14.42 μL aliquots of 10-20 mM ligand solution, it was concluded that there was no binding. The mean enthalpies measured from the injection of effectors into the buffer were subtracted from raw titration data prior to data analysis with the MicroCal version of ORIGIN. Data were fitted with the ‘One binding site model’ of ORIGIN.

**Abbreviations**: TMAO, trimethylamine N-oxide; LBD, Ligand binding domain

**Conflict of interest**: The authors do not declare a conflict of interest.

## Acknowledgements

**Acknowledgements:** This work was supported by the Spanish Ministry for Science and Innovation/*Agencia Estatal de Investigación* 10.13039/501100011033 (grants PID2020-112612GB-I00 to TK and PID2019-103972GA-I00 to MAM) and the Junta de Andalucía (grant P18-FR-1621 to TK) and by the US National Institutes of Health (grant R35GM131760 to IBZ). JPCV was supported by the grant Unión Europea-NextGenerationEU RD 289/2021 UPM-Recualifica Margarita Salas.

## Notes

### Competing Interest Statement

The authors have declared no competing interest.

