## Supplemental figures and tables for "Amine recognizing domain in diverse receptors from bacteria and archaea evolved from the universal amino acid sensor"

This file contains:

- Supplementary figures S1 through S6
- Supplementary tables S1 through S4
- Supplementary references

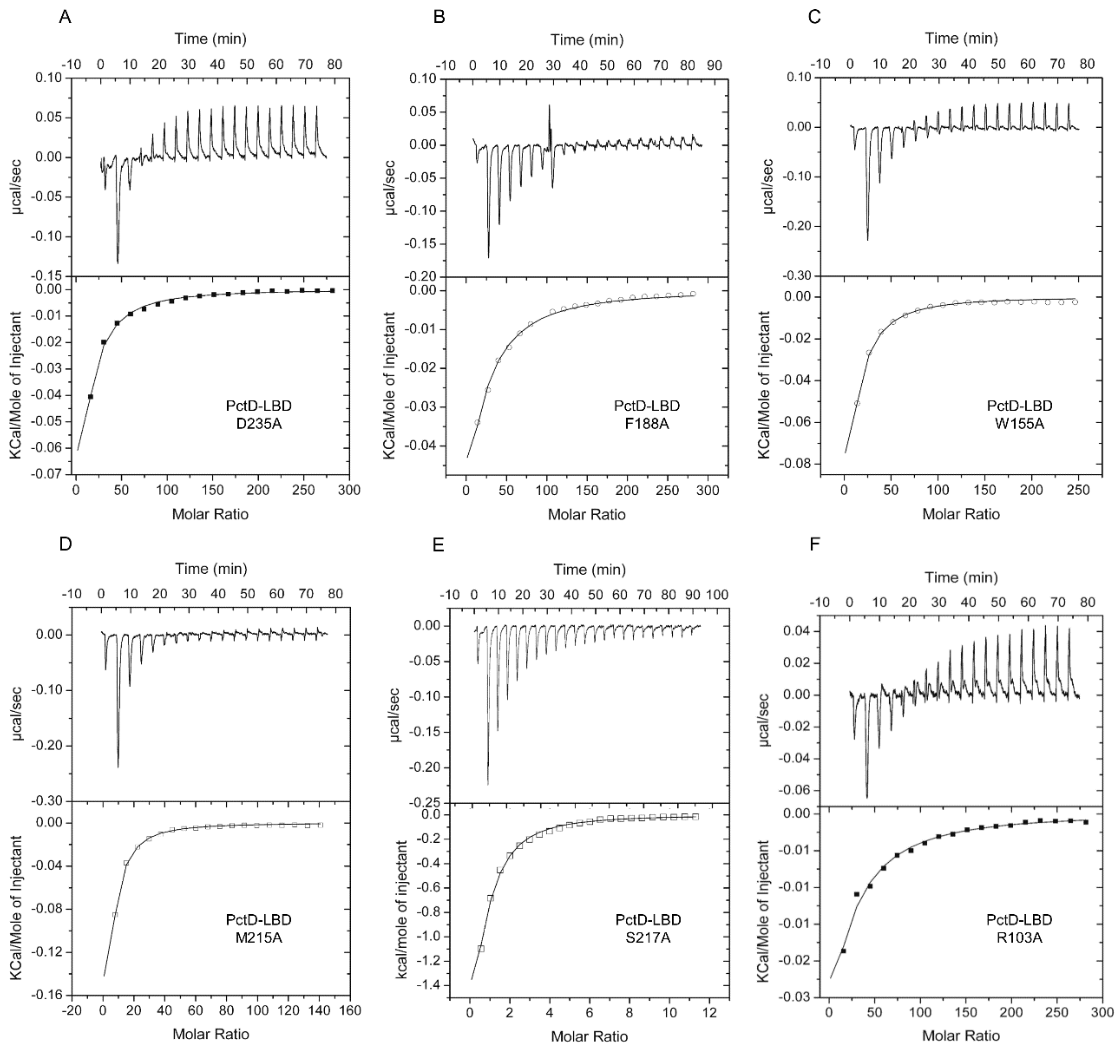


**Fig. S1) Microcalorimetric titrations of PctD-LBD mutants with choline**. Proteins at 14 to 16 µM were placed into the sample cell and titrated with 9.6 to 14.42 µL aliquots of 1mM (PctD-LBD S217A), 10 mM (PctD-LBD M215A) or 20 mM choline (the remaining mutants) solutions made up in dialysis buffer. Upper panels: raw titration data. Lower panels: integrated, concentration-normalized and dilution heat-corrected peak areas and best fit using the "one–binding site model" of the MicroCal version of ORIGIN. The derived thermodynamic parameters are shown in Table. 1.

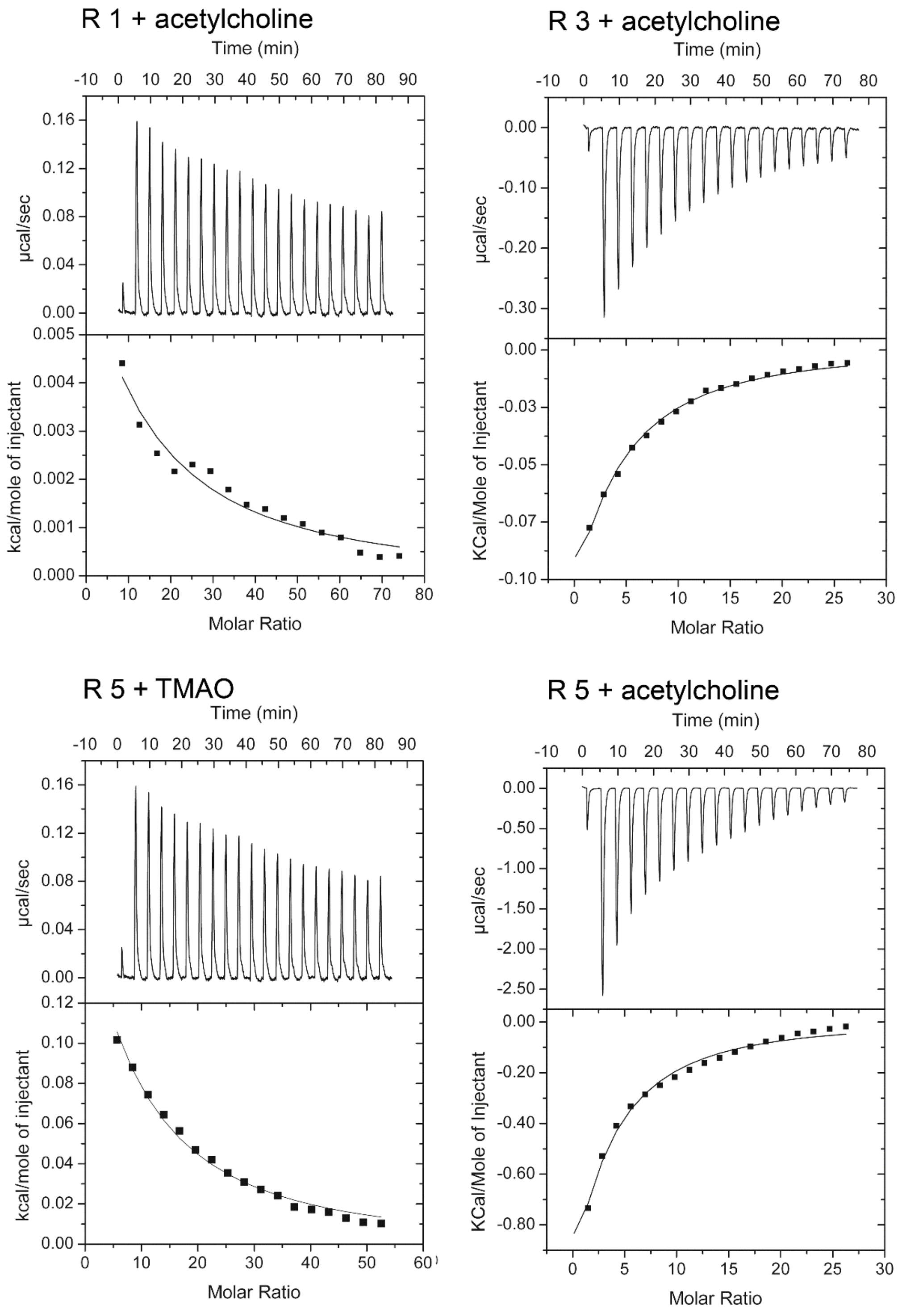


**Fig. S2) Microcalorimetric titrations of predicted amine responsive dCache domains with different quaternary amines.** Upper panel: Raw data for the titration of 75 µM of protein with 14.42 µl aliquots of 10 to 20 mM of quaternary amines. Lower panel: Concentration-normalized and dilution heat corrected integrated raw data. The line is the best fit using the “One binding site model” of the MicroCal version of ORIGIN.

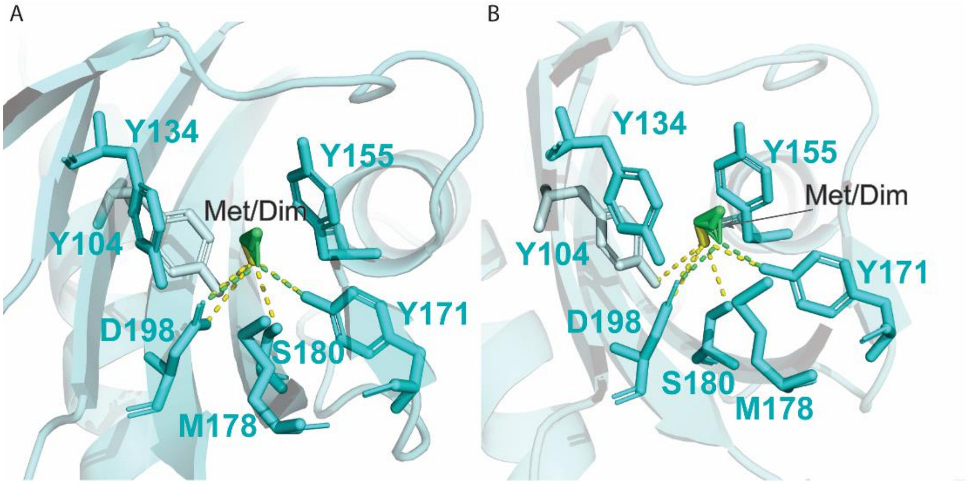


**Fig. S3) Ligand binding module of the dCache_1 domain from the archaeon *Methanosarcina mazei* docked with methylamine (Met, in yellow) and dimethylamine (Dim, in green). A** and **B** are two slightly different angles. Predicted polar contacts are shown.

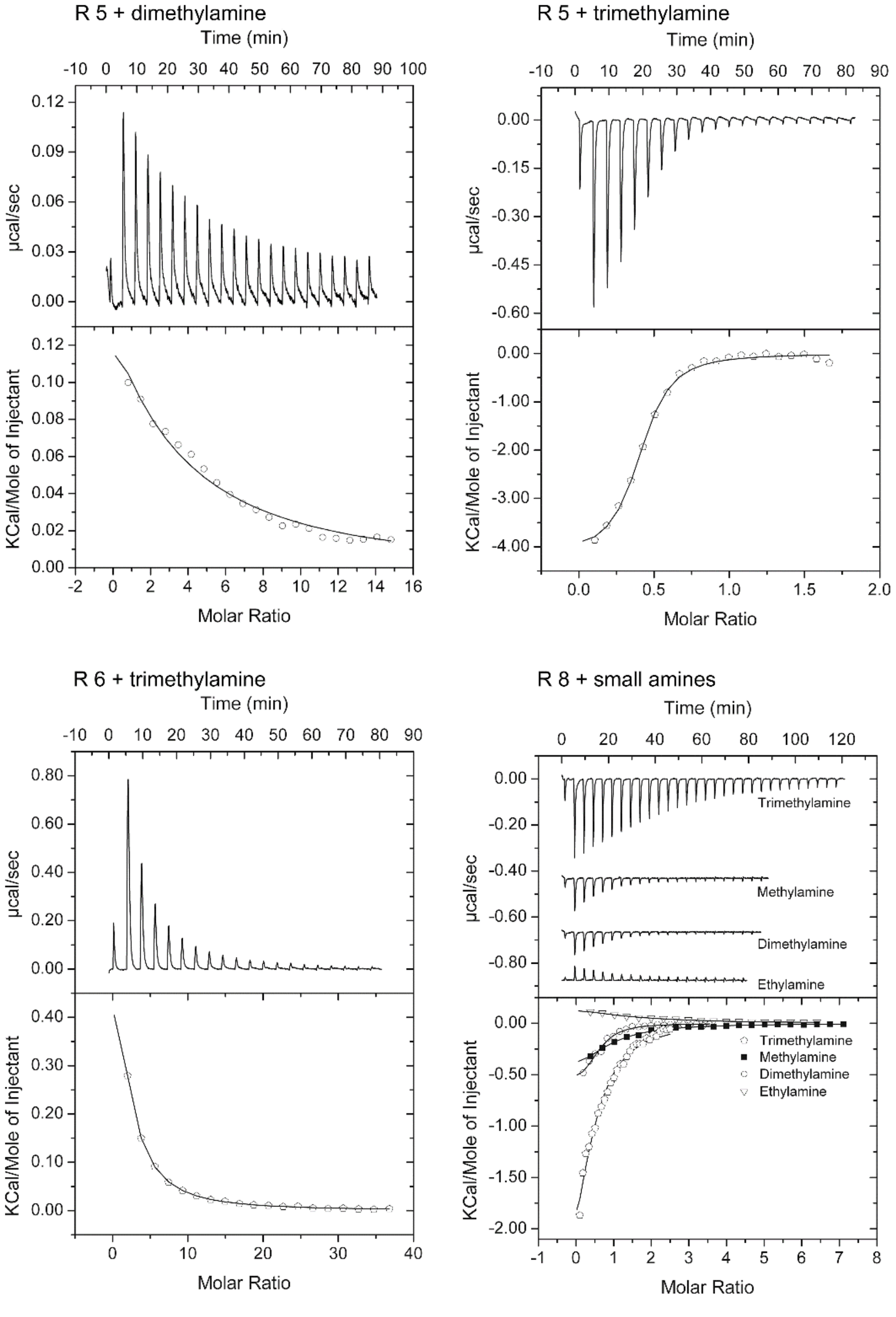


**Fig. S4) Microcalorimetric titrations of predicted amine responsive dCache domains with different small amines.** Upper panel: Raw data for the titration of 30 to 50 µM of protein with 3.2 to 12.8 µl aliquots of 1 to 10 mM of small amines. Lower panel: Concentration-normalized and dilution heat corrected integrated raw data. The line is the best fit using the "One binding site model" of the MicroCal version of ORIGIN.

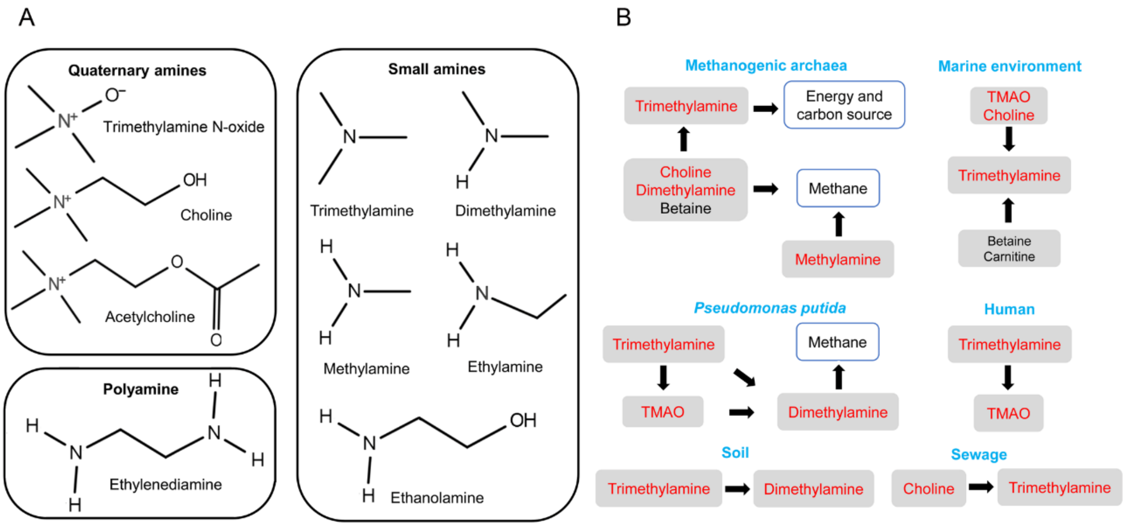


**Fig. S5) Structures of ligands recognized by this domain family (A) and major metabolic processes involving these ligands (B).** Ligands recognized by members of this domain family are shown in red. Information on the metabolic processes have been obtained from the following sources: methanogenic archaea (1, 2), marine environment (3, 4), *Pseudomonas putida* (5), human (6), soil (7) and sewage(8).

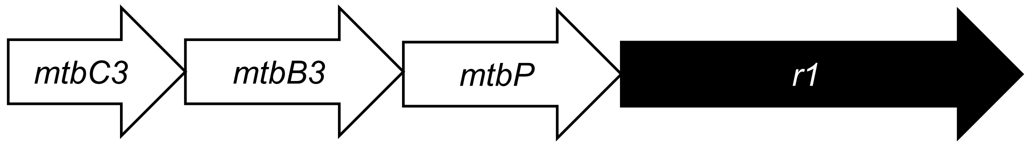


*mtbC3:* Dimethylamine-corrinoid protein
*mtbB3:* Dimethylamine-corrinoid methyltransferase
*mtbP:* APC family permease
*r1:* Receptor 1

**Fig. S6) Genetic environment of the gene encoding R1 (Histidine kinase from *Methanosarcina mazei* S6).** The apo-3D structure of the R1-LBD has been reported (9). There is 100 % sequence identity between the above four genes in *M. mazei* S6 and *M. mazei* Gö1. The latter strain was used in a study the assess the effect of trimethylamine on gene transcript levels, showing large increases of *mtbC3* and *mtbB3* transcript levels in the presence of trimethylamine (10).

**Table S1) Increases in the midpoint of the thermal unfolding transition of protein (ºc) in the presence of a given ligand as compared to ligand-free protein.** Conditions in which binding was observed by isothermal Titration Calorimetry are shown in bold.

| Protein | L-Carnitine | Betaine | L-Pro | Methylamine | Ethylamine | Dimethylamine | Trimethylamine | Ethanolamine | Ethylenediamine |
|---|---|---|---|---|---|---|---|---|---|
| R1 | 0.44 | -0.59 | 0 | **3.74** | **2.41** | **12.5** | **8.5** | -0.18 | -0.24 |
| R3 | -0.35 | -0.78 | -0.35 | 0.2 | 0.1 | 0.5 | 0.75 | **13.9** | **6.9** |
| R4 | 0.20 | 1.67 | 1.00 | 0.59 | 0.6 | 0.9 | 3.84 | 1 | 0.8 |
| R5 | -0.27 | -0.26 | -0.53 | 0.27 | 0.84 | **3.45** | **10.49** | 0.85 | 1.52 |
| R6 | 1.13 | -0.26 | 0.73 | 0.2 | 0.1 | 5 | **4.8** | 0.3 | -0.2 |
| R7 | 0 | -0.04 | -1.83 | **6** | 0.1 | **12** | **9** | **15** | **10** |
| R8 | 0.40 | -0.6 | -1.00 | **5.29** | **3.68** | **6.63** | **7.13** | 0.2 | 0.3 |
| R9 | 0.67 | 0 | 0.67 | 0.2 | 0.3 | -0.1 | -0.3 | 0.1 | 0.2 |
| R10 | 0.27 | -1.22 | 0.27 | 0.19 | -0.21 | -0.71 | -0.48 | 0.09 | 0.23 |
| PacA-LBD | nd | nd | nd | 0.1 | 0.2 | 0.1 | **5** | 0.1 | 0.2 |

**Table S2) Strains and plasmids used in this study.**

| | **Relevant characteristics** | **Reference or source** |
|---|---|---|
| **Strains** | | |
| *Escherichia coli* BL21 (DE3) | $F^-$ *ompT gal dcm lon hsdS*$_B$($r_B^- m_B^-$) λ(DE3 [*lacI lacUV5-T7p07 ind1 sam7 nin5*]) [*malB*$^+$]$_{K-12}$($λ^S$) | (11) |
| *E. coli* BL21-AI | F- *ompT hsdS*$_B$ ($r_B^- m_B^-$) *gal dcm araB::T7RNAP-tetA* | Invitrogen |
| **Plasmids** | | |
| pET28_ECA_RS10935 -LBD | $Km^R$; pET28b(+) derivative containing a DNA fragment encoding PacA-LBD (*Pectobacterium atrosepticum* SCRI1043) | (12) |
| pET28-PctD-LBD-D235A | $Km^R$; pET28b(+) derivative containing DNA fragment encoding PctD-LBD D235A | GenScript |
| pET28- PctD-LBD-F188A | $Km^R$; pET28b(+) derivative containing DNA fragment encoding PctD-LBD F188A | GenScript |
| pET28- PctD-LBD-W155A | $Km^R$; pET28b(+) derivative containing DNA fragment encoding PctD-LBD W155A | GenScript |
| pET28- PctD-LBD-M215A | $Km^R$; pET28b(+) derivative containing DNA fragment encoding PctD-LBD M215A | GenScript |
| pET28- PctD-LBD-S217A | $Km^R$; pET28b(+) derivative containing DNA fragment encoding PctD-LBD S217A | GenScript |
| pET28- PctD-LBD-R103A | $Km^R$; pET28b(+) derivative containing DNA fragment encoding PctD-LBD R103A | GenScript |
| pET28-R1-LBD | $Km^R$; pET28b(+) derivative containing DNA fragment encoding R1-LBD | GenScript |
| pET28-R2-LBD | $Km^R$; pET28b(+) derivative containing DNA fragment encoding R2-LBD | GenScript |
| pET28-R3-LBD | $Km^R$; pET28b(+) derivative containing DNA fragment encoding R3-LBD | GenScript |
| pET28-R4-LBD | $Km^R$; pET28b(+) derivative containing DNA fragment encoding R4-LBD | GenScript |
| pET28-R5-LBD | $Km^R$; pET28b(+) derivative containing DNA fragment encoding R5-LBD | GenScript |
| pET28-R6-LBD | $Km^R$; pET28b(+) derivative containing DNA fragment encoding R6-LBD | GenScript |
| pET28-R7-LBD | $Km^R$; pET28b(+) derivative containing DNA fragment encoding R7-LBD | GenScript |
| pET28-R8-LBD | $Km^R$; pET28b(+) derivative containing DNA fragment encoding R8-LBD | GenScript |
| pET28-R9-LBD | $Km^R$; pET28b(+) derivative containing DNA fragment encoding R9-LBD | GenScript |
| pET28-R10-LBD | $Km^R$; pET28b(+) derivative containing DNA fragment encoding R10-LBD | GenScript |

Km = Kanamycin

**Table S3) Sequences of recombinant proteins used in this study.**

| Protein | Sequence |
|---|---|
| **PacA-LBD** | MGSSHHHHHHSSGLVPRGSHMSWQSSSEQKSLAERYLQQIAQSEALRIQQE<br>LNYARDVAHNLGQGLAALPSAGIKDRAVVDKMMEYALRDNPEYLSISVIFE<br>ENVFDGRDAEFADQPGQAPKGRYAWFVDRDQAGNYAMHPLLSYLTPGQGDY<br>YLLPQKSQKDTLIEPYTYAYNGVPTLLTSVAAPIVSQGKLWGVVTSDISLA<br>SLQQKINQIKPWEGGGYAMLLSSAGKVISYPDKSQTSKAWQGPTDNFTSSV<br>VQHDDAILGEQALVTWQPVTIGNSTEKWYLGIVVPVSQVMAASERQL |
| **PctD-LBD R103A** | MGSSHHHHHHSSGLVPRGSHMAGARTQELVQQRTQGLLEKVINERLVALAR<br>AQVSQIQRELEYPLTVVHGLANSTRLLGEPGADGMPQLNASADEISALLRS<br>TVQNNPKLLDTFMAWEPNAFDTDAAFAGQPGKGYGPDGRYLPWWYRGADGK<br>PIVEAMADSIDSEKLLPTGVRENEFYACPKENKRPCIIDPAPYEMGGKTVM<br>MSSFNVPIMVGDQFRGAVGADLSLAFIQDLLKRADQQLYDGAGEMALIASN<br>GRLVAYTRDDSKLGEPAGSVLDGNEVDNLKNLTVDQPLYDIDAEHGHIELF<br>LPFTIADSGVRWTLMLQIPQAAVFGELQQLQGELSDQRQQDILGM |
| **PctD-LBD W155A** | MGSSHHHHHHSSGLVPRGSHMAGARTQELVQQRTQGLLEKVINERLVALAR<br>AQVSQIQRELEYPLTVVHGLANSTRLLGEPGADGMPQLNASRDEISALLRS<br>TVQNNPKLLDTFMAWEPNAFDTDAAFAGQPGKGYGPDGRYLPAWYRGADGK<br>PIVEAMADSIDSEKLLPTGVRENEFYACPKENKRPCIIDPAPYEMGGKTVM<br>MSSFNVPIMVGDQFRGAVGADLSLAFIQDLLKRADQQLYDGAGEMALIASN<br>GRLVAYTRDDSKLGEPAGSVLDGNEVDNLKNLTVDQPLYDIDAEHGHIELF<br>LPFTIADSGVRWTLMLQIPQAAVFGELQQLQGELSDQRQQDILGM |
| **PctD-LBD F188A** | MGSSHHHHHHSSGLVPRGSHMAGARTQELVQQRTQGLLEKVINERLVALAR<br>AQVSQIQRELEYPLTVVHGLANSTRLLGEPGADGMPQLNASRDEISALLRS<br>TVQNNPKLLDTFMAWEPNAFDTDAAFAGQPGKGYGPDGRYLPWWYRGADGK<br>PIVEAMADSIDSEKLLPTGVRENEAYACPKENKRPCIIDPAPYEMGGKTVM<br>MSSFNVPIMVGDQFRGAVGADLSLAFIQDLLKRADQQLYDGAGEMALIASN<br>GRLVAYTRDDSKLGEPAGSVLDGNEVDNLKNLTVDQPLYDIDAEHGHIELF<br>LPFTIADSGVRWTLMLQIPQAAVFGELQQLQGELSDQRQQDILGM |
| **PctD-LBD M215A** | MGSSHHHHHHSSGLVPRGSHMAGARTQELVQQRTQGLLEKVINERLVALAR<br>AQVSQIQRELEYPLTVVHGLANSTRLLGEPGADGMPQLNASRDEISALLRS<br>TVQNNPKLLDTFMAWEPNAFDTDAAFAGQPGKGYGPDGRYLPWWYRGADGK<br>PIVEAMADSIDSEKLLPTGVRENEFYACPKENKRPCIIDPAPYEMGGKTVM<br>ASSFNVPIMVGDQFRGAVGADLSLAFIQDLLKRADQQLYDGAGEMALIASN<br>GRLVAYTRDDSKLGEPAGSVLDGNEVDNLKNLTVDQPLYDIDAEHGHIELF<br>LPFTIADSGVRWTLMLQIPQAAVFGELQQLQGELSDQRQQDILGM |
| **PctD-LBD S217A** | MGSSHHHHHHSSGLVPRGSHMAGARTQELVQQRTQGLLEKVINERLVALAR<br>AQVSQIQRELEYPLTVVHGLANSTRLLGEPGADGMPQLNASRDEISALLRS<br>TVQNNPKLLDTFMAWEPNAFDTDAAFAGQPGKGYGPDGRYLPWWYRGADGK<br>PIVEAMADSIDSEKLLPTGVRENEFYACPKENKRPCIIDPAPYEMGGKTVM<br>MSAFNVPIMVGDQFRGAVGADLSLAFIQDLLKRADQQLYDGAGEMALIASN<br>GRLVAYTRDDSKLGEPAGSVLDGNEVDNLKNLTVDQPLYDIDAEHGHIELF<br>LPFTIADSGVRWTLMLQIPQAAVFGELQQLQGELSDQRQQDILGM |
| **PctD-LBD D235A** | MGSSHHHHHHSSGLVPRGSHMAGARTQELVQQRTQGLLEKVINERLVALAR<br>AQVSQIQRELEYPLTVVHGLANSTRLLGEPGADGMPQLNASRDEISALLRS<br>TVQNNPKLLDTFMAWEPNAFDTDAAFAGQPGKGYGPDGRYLPWWYRGADGK<br>PIVEAMADSIDSEKLLPTGVRENEFYACPKENKRPCIIDPAPYEMGGKTVM<br>MSSFNVPIMVGDQFRGAVGAALSLAFIQDLLKRADQQLYDGAGEMALIASN<br>GRLVAYTRDDSKLGEPAGSVLDGNEVDNLKNLTVDQPLYDIDAEHGHIELF<br>LPFTIADSGVRWTLMLQIPQAAVFGELQQLQGELSDQRQQDILGM |
| **R1** | MGSSHHHHHHSSGLVPRGSHMTTQEEKLAYQQSVEMASNYANQFDADMKAN<br>LAIARTISTTMESYETADRDEALLILENLLRDNPHLLGTYVAFEPDAFDGK<br>DAEYTNSPAHDGTGRFVPYWNKMNGTASVAPLLHYDSSDYYQLPKATEKDV<br>LTEPYFYEGVFMVSYVSPIMKEGEFAGIGGVDVSLEYVDEVVSKVRTFDTG |

| | |
|---|---|
| | YAFMVSNSGVILSHPTHKDWIGKKDLYDFGGEELEKASRDIKNGIGGHLET<br>ADPTTGKTVILFYEPVETGDFAFVLVVPKEEMLAGVADLRER |
| **R2** | MGSSHHHHHHSSGLVPRGSHMQLMKLYDVSLRQGELVAQNQSNAYTTKMSI<br>ETNDALIRLEGLQQSLQQMKEYNMTDRSEAVRLIENFVREQPYILGVFTVW<br>EPNAFDNQDGNFRNKSSYDDDTGRFVPYIVRQGDKIVAYPNKNYENIGDGD<br>YYQIPKRTKKFALMEPYYYDINGERILISSFVYPILDEQGKFLGVVGADIS<br>LDMVQQEVEKIRPMGGYATMITAGDSYLANGFDRALVSKPYLPLPKGESLE<br>ELKEQALTIMYTSDPMLGGTVMRLLNPIHIKDQTWYFETIIPKGNMLKDYY<br>KGLSNT |
| **R3** | MGSSHHHHHHSSGLVPRGSHMHTSSKTTLMQEARADAANLTLASIRKIEGT<br>LASVEAIPGLLAFSYGKNKPTASAISTDLLGFILFNSAVYGSCVAYEPYAF<br>DRDVEFFAPYAYMPGGRPMFTYLSADYNYPQADWFLIPKEIRRPIWSEPYF<br>DEGGGNVVMSTYSIPFFREEDGRKRFLGVVTADISLEWLRTFIKSISIYQS<br>GYAFLLSRNGVFLSHPNDQFIMRESIFSLAETHSSKVLRDIGKKMVQGETG<br>FVRLPEFVMGEPAWLSYAPVSNSDWSMGLVIPEAEMFQGLEGLSRE |
| **R4** | MGSSHHHHHHSSGLVPRGSHMSNRSIEMAQKDAFSLAQETADKYKNAIIAE<br>LQGARITAETFSTVFETLKDYNLTDRDMMNDILKNALANKEYITAFCIAYD<br>PNAMDGKDAQYAGQGPAYDETGRYAPYWNKLGGNIDVEFLPDIDSEDWYIV<br>PKAERHEYITDPYPYGLQGRTVMLASLIFPIIHSDKFIGIISSDIVLDKLQ<br>EMVDKVNPHGQEGYTEIISHSGAVIAHPNKDYLGKDLEETLVEGQSRLQHI<br>DEIKSAINSGEMYISTGKNFYTVYMPIQFSSVTNPWSVAVSIPMAKILANA<br>DSIRNY |
| **R5** | MGSSHHHHHHSSGLVPRGSHMTYNSLKTATVSSTEISNQMAVTYANQVVDK<br>MDDAMSAARSLAHALSGVIGKNVSRQAIQQMAGSILLGDEDFLGYTVCFEP<br>NAYDAKDAFFANKPGHDNTGRFVSYMTKNGSGGFVVEPLVDYENESAAPWY<br>WIPMRTMKEFVTEPLMYPIQGKNVYMVSFMCPIITNGKFVGVTGVDLSINY<br>LQDMVVKANVFDGHGNFDIVSHQGVFAANSGNPDFVGKNILEQKNIGAEDQ<br>LVDIEKGNSLTRIDNGILKAFVPVIVGRCPTAWQVSISVPVDYITQEARAQ<br>MIYQ |
| **R6** | MGSSHHHHHHSSGLVPRGSHMRVVHSARQEANALSRTKAQAIGAEMAHRLG<br>RAIGTARTLSEALEGILAEGHPSRAQADAMLRGSLEGNTDYIGVWTLWEPN<br>AFDGRDADYVNKPGHDATGRYIAYWNRGSGKVIVEPLVDYTTEGAGDYYLL<br>AKHSNQETVLEPYIYKVAGRDVLMTSLVVPVNRADGTFAGVVGVDLPLETL<br>GAEIAKVKVGETGYAALVSNTGIYAAHPRAERLGKPMKDTDPWVVPFLGNL<br>KKGEAFETESFSRTLNDMTYRFGVPVRIGSSSTPWCVSITIRESEVLAGAW<br>KLRNT |
| **R7** | MGSSHHHHHHSSGLVPRGSHMSYINARNEALNAAQKRAQIVAKNYSKEISD<br>ELGQAITVAENLGSMLKGRIRNENATLTRDEVSEIFKNALADNPQFIGISI<br>AFEPNAFDSLDAQFDGDSRYYEKGQFATYFVRGNLNSGKDMYSTITQEPLR<br>DLEISDYYIVPKQTLSNVMIEPYIYEVQGKEVLMTTCSSPIVIDGKYYGVV<br>GVDIEVDFIKELVKGSENDMEFKDILIISSKGNIVGSKYEMAYNQDENLKE<br>EILHFKQGSHVEYSNGIFDVYELINIRDIDDKWGIKLSVDQKTIMGSASRL<br>LANQ |
| **R8** | MGSSHHHHHHSSGLVPRGSHMTTKSGSDIETLAFQSGEQLGHRYGEMVHAR<br>LGNAMEAGRFIATSLVGLKAAGRTDREQLSIWLKSIAEANPDFLGVWVGME<br>PNALDGRDAEFANKPGSDASGRFLPYWNRGSGTVALESLVGYDEPGSDGAY<br>YQIPKRTGHAMVVEPYSYTVAGRKVLMVSMSVPIVENGRVIGVAGIDLSTD<br>GIWSMLKTVKPFDSGSIHLISNDGVWAGHPDSERMGQPIGKSDPALDAAKP<br>AIRAGRSFEQMSVADGQPVKQLFLPVTVAGTETPWSLLVNLPLDKINAPVR<br>ELRNAT |
| **R9** | MGSSHHHHHHSSGLVPRGSHMYNARASQQTAKLQSSESVIDKSQQLLQTGA<br>LLNATEISEYLSEAIYRAEMLAANALFLKNNSEENFGESEALRTSLDEMVR<br>KSVLGFDTIEGAYLVFRPNMLDSEDSNYVNADYVGSNDIGQFAAYWTKAAN<br>GQNVISRVLTQAQLTEESNKERFVCPIEQASPCITSPRMVEFETERYLATS<br>LSVPILIDGVAIGFYGIDLTLAPLIGITQKSDNNLFDGQGKVSIVSENNAL |

| | |
|---|---|
| | VASDADFLTLGETFQSENLSRSTVSSLLQAGQVNTQWSEDGQWLVVFAPTK<br>VANQNWGVIFEMPRQSVMQDAEQLDILLTEQLERGIRSE |
| **R10** | MGSSHHHHHHSSGLVPRGSHMTTLVYNNDKKSALLYMESLAAEKANIAKLE<br>METALETARTLASVFSTWENIPVEERRTLFSGILKTVVEKNEDFQGAWTCW<br>EENTLDASDSFYKGLPGYDETGRFIPYWYRSDSGRIEYEPLTGYTQPGEGN<br>YYLVPLNNKKEAAAEPYIYELKGKPRWLTSLSVPIYDNANRVAGIVGINLS<br>LDHLQSHLSDLVFFDTGFGRLVSAEGLVVTHPDRDRIGKIIGEFVKDTGQA<br>LINSIKGGEATSGEAWSESLESMTTKTNVPFSIGRTETNWFYGTVVPSHEL<br>YANALGFAK |

**Table S4) Buffers used in the purification and analysis of proteins.**

| Prot. | Purification buffer A | Purification buffer B | Analysis buffer |
|---|---|---|---|
| R1 | 30 mM Tris, 300 mM NaCl, 5 % glycerol (vol/vol), 10 mM imidazole, pH 7.5 | 30 mM Tris, 300 mM NaCl, 5 % glycerol (vol/vol), 500 mM imidazole, pH 7.5 | 5 mM Tris, 5 mM MES, 5 mM PIPES, 150 mM NaCl, 10% (vol/vol) glycerol, pH 7.5 |
| R2 | 30 mM Tris, 300 mM NaCl, 5 % glycerol (vol/vol), 10 mM imidazole, pH 7.5 | 30 mM Tris, 300 mM NaCl, 5 % glycerol (vol/vol), 500 mM imidazole, pH 7.5 | 5 mM Tris, 5 mM MES, 5 mM PIPES, 150 mM NaCl, 10% (vol/vol) glycerol, pH 7.5 |
| R3 | 30 mM Tris, 300 mM NaCl, 5 % glycerol (vol/vol), 10 mM imidazole, 0.1 mM EDTA, 5 mM β-mercaptoethanol, pH 7.5 | 30 mM Tris, 300 mM NaCl, 5 % glycerol (vol/vol), 500 mM imidazole, 0.1 mM EDTA, 5 mM β-mercaptoethanol, pH 7.5 | 5 mM Tris, 5 mM MES, 5 mM PIPES, 150 mM NaCl, 10% (vol/vol) glycerol, 0.1 mM EDTA, 5 mM β-mercaptoethanol, pH 8.0 |
| R4 | 30 mM Tris, 300 mM NaCl, 5 % glycerol (vol/vol), 10 mM imidazole, 0.1 mM EDTA, 5 mM β-mercaptoethanol, pH 7.5 | 30 mM Tris, 300 mM NaCl, 5 % glycerol (vol/vol), 500 mM imidazole, 0.1 mM EDTA, 5 mM β-mercaptoethanol, pH 7.5 | 5 mM Tris, 5 mM MES, 5 mM PIPES, 150 mM NaCl, 10% (vol/vol) glycerol, 0.1 mM EDTA, 5 mM β-mercaptoethanol, pH 8.0 |
| R5 | 30 mM Tris, 300 mM NaCl, 5 % glycerol (vol/vol), 10 mM imidazole, 0.1 mM EDTA, 5 mM β-mercaptoethanol, pH 7.5 | 30 mM Tris, 300 mM NaCl, 5 % glycerol (vol/vol), 500 mM imidazole, 0.1 mM EDTA, 5 mM β-mercaptoethanol, pH 7.5 | 5 mM Tris, 5 mM MES, 5 mM PIPES, 150 mM NaCl, 10% (vol/vol) glycerol, 0.1 mM EDTA, 5 mM β-mercaptoethanol, pH 8.0 |
| R6 | 30 mM Tris, 300 mM NaCl, 5 % glycerol (vol/vol), 10 mM imidazole, 0.1 mM EDTA, 5 mM β-mercaptoethanol, pH 7.5 | 30 mM Tris, 300 mM NaCl, 5 % glycerol (vol/vol), 500 mM imidazole, 0.1 mM EDTA, 5 mM β-mercaptoethanol, pH 7.5 | 5 mM Tris, 5 mM MES, 5 mM PIPES, 150 mM NaCl, 10% (vol/vol) glycerol, 0.1 mM EDTA, 5 mM β-mercaptoethanol, pH 8.0 |
| R7 | 30 mM Tris, 300 mM NaCl, 5 % glycerol (vol/vol), 10 mM imidazole, 0.1 mM EDTA, 5 mM β-mercaptoethanol, pH 7.5 | 30 mM Tris, 300 mM NaCl, 5 % glycerol (vol/vol), 500 mM imidazole, 0.1 mM EDTA, 5 mM β-mercaptoethanol, pH 7.5 | 5 mM Tris, 5 mM MES, 5 mM PIPES, 150 mM NaCl, 10% (vol/vol) glycerol, 0.1 mM EDTA, 5 mM β-mercaptoethanol, pH 8.0 |
| R8 | 30 mM Tris, 300 mM NaCl, 5 % glycerol (vol/vol), 10 mM imidazole, pH 7.5 | 30 mM Tris, 300 mM NaCl, 5 % glycerol (vol/vol), 500 mM imidazole, pH 7.5 | 5 mM Tris, 5 mM MES, 5 mM PIPES, 150 mM NaCl, 10% (vol/vol) glycerol, pH 7.5 |
| R9 | 30 mM Tris, 300 mM NaCl, 5 % glycerol (vol/vol), 10 mM imidazole, pH 7.5 | 30 mM Tris, 300 mM NaCl, 5 % glycerol (vol/vol), 500 mM imidazole, pH 7.5 | 5 mM Tris, 5 mM MES, 5 mM PIPES, 150 mM NaCl, 10% (vol/vol) glycerol, pH 7.5 |

| | | | |
|---|---|---|---|
| R10 | 30 mM Tris, 300 mM NaCl, 5 % glycerol (vol/vol), 10 mM imidazole, 0.1 mM EDTA, 5 mM β-mercaptoethanol, pH 7.5 | 30 mM Tris, 300 mM NaCl, 5 % glycerol (vol/vol), 500 mM imidazole, 0.1 mM EDTA, 5 mM β-mercaptoethanol, pH 7.5 | 5 mM Tris, 5 mM MES, 5 mM PIPES, 150 mM NaCl, 10% (vol/vol) glycerol, pH 7.5 |
